# Cytokinin oxidase/dehydrogenase family genes exhibit functional divergence and overlap in rice growth and development, especially in control of tillering

**DOI:** 10.1101/2021.05.09.443313

**Authors:** Chenyu Rong, Yuexin Liu, Zhongyuan Chang, Ziyu Liu, Yanfeng Ding, Chengqiang Ding

## Abstract

Cytokinins play key roles in plant growth and development; hence, cytokinin biosynthesis and degradation have been extensively studied. Cytokinin oxidase/dehydrogenases (CKXs) are a group of enzymes that regulate oxidative cleavage to maintain cytokinin homeostasis. In rice, 11 *OsCKX* genes have been identified to date; however, most of their functions remain unknown. Here, we comprehensively analyzed the expression patterns and functions of *OsCKX* genes. Using CRISPR/Cas9 technology, we constructed mutants of all *OsCKX* genes to determine the functions of OsCKXs in rice development. The results revealed that the single *osckx* and higher-order *osckx4 osckx9* mutant lines showed functional overlap and subfunctionalization. Notably, the *osckx1 osckx2* and *osckx4 osckx9* double mutants displayed contrasting phenotypic changes in tiller number and panicle size compared to the wild type. Moreover, we identified several genes with significantly altered expression in *osckx4* and *osckx9* single and double mutant plants. Many differentially expressed genes were found to be associated with auxin and cytokinin pathways. Additionally, the cytokinins in *osckx4 osckx9* mutants were increased compared to the wild type. Overall, our findings provide new insights into the functions of *OsCKX* genes in rice growth and may be used as a foundation for future studies aimed at improving rice yield. Key words: Cytokinin, expression pattern, *OsCKX*, panicle, phenotype, rice, tiller

**Highlight:** The osckx4 osckx9 double mutant had a significantly greater number of tillers, whereas the osckx1 osckx2 double mutant showed the opposite phenotypic change, compared to the wild type

## INTRODUCTION

Cytokinin controls plant growth and development by promoting plant cell division, growth, and differentiation. Specifically, it is responsible for stem enlargement and differentiation, organ development and structure, nutrition absorption, senescence, and stress response (Zurcher *et al*., 2016). Cytokinin biosynthesis starts from the ADP, ATP, or tRNA catalyzed by different adenosine phosphate isopentenyl transferases and activated by LONELY GUY enzymes (Kurakawa *et al*., 2007). After performing its function, cytokinin is inactivated by certain enzymes such as cytokinin oxidase/dehydrogenases (CKXs) that induce cytokinin oxidative cleavage, which is an irreversible process. CKXs are essential for maintaining cytokinin homeostasis during plant growth and development. Therefore, extensive research on CKXs has been conducted and is still ongoing. Each plant species has a group of *CKX* genes: for example, *Arabidopsis* has seven, rice has 11, and maize has 13 (Ashikari *et al*., 2005; Zalabak *et al*., 2016). Most CKXs in maize and *Arabidopsis* have putative N-terminal secretory peptides that assist in locating the endoplasmic reticulum (ER) (Niemann *et al*., 2018; Zalabak *et al*., 2016). Previous studies have discovered several functions of CKXs in *Arabidopsis* and other crops. The overexpression of *AtCKX1* and *AtCKX2* in tobacco resulted in smaller shoots, larger roots, and smaller shoot apical meristems (SAMs), while *AtCKX7* overexpression in *Arabidopsis* resulted in shorter primary roots (Kollmer *et al*., 2014; Werner *et al*., 2001). Furthermore, *ckx3 ckx5* mutants developed larger SAM and more siliques in *Arabidopsis* (Bartrina *et al*., 2011). Sextuple *ckx3 ckx5* mutants of oilseed rape showed larger and more active inflorescence meristems (Schwarz *et al*., 2020). In barley, *HvCKX1* downregulation resulted in more spikes, more grains per spike, and a higher 1,000-grain weight (Zalewski *et al*., 2010). *TaCKX6-D1* was found to be associated with grain filling and grain size in wheat (Zhang *et al*., 2012).

In rice, *OsCKX2/Gn1a* was the first *CKX* gene identified. Rice varieties with low *OsCKX2* expression yielded more grains per panicle, while *OsCKX2* downregulation or *osckx2* mutation produced more tillers, more grains per panicle, and heavier grains (Ashikari *et al*., 2005; Yeh *et al*., 2015). *OsCKX4* overexpression reduced the tiller number, grain number per panicle, grain weight, and plant height, while it increased the number of roots (Gao *et al*., 2014). On the other hand, *osckx9* mutants developed more tillers and smaller panicles, while *OsCKX9* overexpression mutants displayed more tillers, shorter culms, smaller panicles, and lower setting rates (Duan *et al*., 2019). *OsCKX11* was found to regulate leaf senescence and grain number, while *osckx11* mutation resulted in the increased number of spikelets per panicle and tillers (Zhang *et al*., 2021).

In rice breeding, *OsCKX2* has been artificially selected from wild rice since ancient times and can be divided into alleles from different geographic distributions (Ashikari *et al*., 2005; Wang *et al*., 2015). Since most plants with downregulated *CKX* expression exhibit better phenotypes, researchers believed that *CKX* is a potential target to improve yield or to initiate another “Green Revolution” (Ashikari *et al*., 2005; Chen *et al*., 2020; Jameson and Song, 2020). Numerous studies have analyzed the structures of and genetic relationships between *OsCKXs* (Ashikari *et al*., 2005; Zhang *et al*., 2021). However, the expression patterns that may determine their functions during plant growth, especially of *OsCKXs* with similar amino acid sequences, are yet to be elucidated.

*OsCKX* genes play an important role in the crosstalk between cytokinin and other hormones by regulating cytokinin content in plant tissues. Most *OsCKX* genes are upregulated by exogenous cytokinins, such as *trans*-zeatin (tZ), *N*^6^-(Δ ^2^-isopentenyl) adenine (iP) and 6-benzylaminopurine (6-BA) (Duan *et al*., 2019). *OsCKX4* can be induced by auxin, which plays a role in regulating the root system (Gao *et al*., 2014). In addition, *OsCKX9* acts downstream of strigolactones (SLs) and plays a key role in the crosstalk between cytokinin and SLs. The SL-induced activation of *OsCKX9* relies on D53, which acts as a repressor of SL signaling and consequently functions in the SL-induced regulation of tiller development (Duan *et al*., 2019). On the other hand, *OsCKX11* has antagonistic roles between the cytokinin and abscisic acid (ABA) pathways during leaf senescence. In *osckx11* mutants, the downregulation of ABA biosynthesis genes and the upregulation of ABA degradation genes result in the reduction of ABA content in flag leaves and subsequently regulate leaf senescence, which shows the relationship between cytokinin and ABA (Zhang *et al*., 2021). Thus, these findings demonstrate that *OsCKX* genes act as bridges between cytokinin and other plant hormones.

In this study, we investigated the genetic relationships and expression patterns of *OsCKX* genes using RNA sequencing (RNA-seq). We evaluated the phenotypes of *osckx* single and double mutants produced using CRISPR/Cas9 technology to determine the function of different *OsCKX*s. We also analyzed the transcriptomes of the roots and shoot bases from Nipponbare (NIP), *osckx4*, *osckx9*, and *osckx4 osckx9* mutants to identify the genes that were influenced by *OsCKX4* and *OsCKX9*. Our findings may provide new insights regarding the potential application of *OsCKX* genes for improving agricultural traits.

## MATERIALS AND METHODS

### Plant materials and growth conditions

*Oryza sativa* L. ssp. *japonica* (cultivars: Nipponbare (NIP) and Zhonghua 11 (ZH11)) was the wild-type (WT) rice material chosen for this study. The mutants included *osckx3*, *osckx4*, *osckx5*, *osckx1 osckx2*, *osckx9*, *osckx4 osckx9*, and *osckx6 osckx7 osckx10* from NIP and *osckx1*, *osckx2*, *osckx7*, *osckx8*, *osckx9*, and *osckx11* from ZH11 (Supplemental Figure S1–S3). The target for *osckx1 osckx2* matches *OsCKX1* and *OsCKX2* completely, and has a 1-bp mismatch for *OsCKX11*. The sequencing results showed that the *osckx1 osckx2-19* and *osckx1 osckx2-21* line had a 1-bp insertion in *OsCKX1* and *OsCKX2*, respectively, but no change was observed in *OsCKX11* (Supplemental Figure S2A).

The rice plants were grown in a field in Danyang, Jiangsu Province, China (31.907° N 119.466° E). We used 300 kg/hm^2^ nitrogen, 150 kg/hm^2^ P2O5, and 240 kg/hm^2^ K2O as base fertilizers for the field experiment. For the pot experiment, 4.28 g of urea was added to each pot, while 1.92 g of KH2PO4 and 1.49 g KCl were used during the whole growth period. For the hydroponic experiment, the nutrient solution was composed of 2 mM KNO3 and 2 mM NH4Cl, and the concentrations of other nutrient elements were the same as those described by Wang *et al*., 2020. The plants were grown in climate chambers under long-day conditions (15 h light at 28 °C and 9 h dark at 24 °C) with a relative humidity of ∼70%, and the treatment lasted 20 days.

### Plasmid construction for genetic transformation

To construct the mutants for research by using CRISPR/Cas9 technology, as previously described, each *OsCKX* gene was assigned one to two single guide RNA oligo targets by using CRISPR/Cas9 technology as previously described (Mao *et al*., 2013). The primers used for vector construction and genotyping are shown in Supplemental Table S1.

To construct *pOsCKX::GUS*, an upstream fragment of the *OsCKX*-encoding region (>3 kb) was amplified by PCR, and the resulting amplicon was excised with the corresponding restriction endonuclease and ligated into the *pCAMBIA1300::GUS* vector (Wang *et al*., 2020). The primers used for vector construction are shown in Supplemental Table S1. Analysis of GUS activity The GUS reporter activity was assayed by histochemical staining using GUS Staining Kit (FCNCS, https://www.fcncs.com). Various tissues were collected from the *pOsCKX::GUS* transgenic plants at each developmental stage, after which they were immersed in the GUS staining solution and incubated for 6–48 h at 37 °C in the dark. The samples were de-stained thrice with 70% ethanol in a water bath for 5 min. Images were taken using SZX16 microscope (Olympus, Tokyo, Japan; https://www.olympus-lifescience.com/en/). Each *pOsCKX::GUS* transgenic plant had 20 independent lines, from which 2 or 3 independent lines were chosen and subjected to staining at the same position as other lines for further studies. Sampling, RNA extraction, and gene expression analysis

We collected roots from approximately 2 cm under the tip and shoot bases (∼5-mm segments from the first nodes) containing the SAM, axillary buds, young leaves, and tiller nodes from the WT, *osckx4*, *osckx9*, and *osckx4 osckx9* plants at 46 days after sowing. We also collect axillary buds (0-1 cm) in 2021 at 50 days after sowing. The samples were stored at −80 °C until further use.

Total RNA was extracted using an E.Z.N.A.^®^ Plant RNA Kit (Omega Bio-tek Inc., Norcross, GA, USA; https://www.omegabiotek.com/) and subjected to reverse-transcription using a PrimeScript^™^ RT Reagent Kit (Takara Biotechnology, Tokyo, Japan; https://www.takarabio.com/). Quantitative reverse-transcription PCR was performed on an ABI PRISM 7300 Real-Time PCR System (Applied

Biosystems, Thermo Fisher Scientific, Waltham, MA, USA; https://www.thermofisher.com/) with SYBR^®^ Premix Ex Taq^™^ (Takara), following the manufacturer’s instructions. Relative expression analysis was performed using the actin gene as internal control. The primers used are listed in Supplemental Table S1.

### Sequence alignment and phylogenetic analysis

The amino acid sequence alignment was performed using MUSCLE (Madeira *et al*., 2019), and phylogenetic trees were generated using a neighbor-joining method and 1,000 bootstrap iterations in MEGA X (Kumar *et al*., 2018). RNA-Seq and data analysis Before library preparation, Dynabeads® mRNA Purification Kit (Invitrogen, Thermo Fisher Scientific, Waltham, MA, USA; https://www.thermofisher.com/) were used to purify the mRNA. An MGIEasy RNA Library preparation kit was used for library construction from purified mRNA according to the manufacturer’s instruction (BGI, Shenzhen, China; https://www.mgi-tech.com/). The quality of the constructed library was checked and sequenced after passing. High-throughput sequencing was paired-end sequenced on the MGISEQ-2000 platform (BGI). The obtained reads were processed and analyzed, and genes with *Q* values of < 0.05 and log2(fold change) values of > 1 or < -1 were considered to be significantly differentially expressed. Based on the list of DEGs, we created and modified Venn diagrams in Microsoft Excel. Furthermore, we performed KEGG enrichment analysis (*Q* < 0.05) (Kanehisa, 2019), and generated a bubble chart using R.

### Sampling and quantification of cytokinins

From the WT and *osckx4 osckx9* plants, we collected roots from approximately 5 cm below the tip and shoot bases (∼5-mm segments from the first nodes) containing the SAM, axillary buds, young leaves, and tiller nodes at 4 weeks after sowing. The samples were stored at −80 °C until further use. Endogenous cytokinin levels were quantified by Webiolotech (Nanjing, China; https://www.webiolotech.com/) with the ESI-HPLC-MS/MS method, using isotope internal standard as previously reported (Cai et al., 2013).

## ACCESSION NUMBERS

Sequence data from this article can be found in Rice Annotation Project database (http://rice.plantbiology.msu.edu) under the following accession numbers: *Actin* (Os03g0718100), *OsCKX1* (Os01g0187600), *OsCKX2* (Os01g0197700), *OsCKX3* (Os10g0483500), *OsCKX4* (Os01g0940000), *OsCKX5* (Os01g0775400), *OsCKX6* (Os02g0220000), *OsCKX7* (Os02g0220100), *OsCKX8* (Os04g0523500), *OsCKX9* (Os05g0374200), *OsCKX10* (Os06g0572300), *OsCKX11* (Os08g0460600), *OsMADS25* (Os04g0304400), *OsMADS57* (Os02g0731200), *OsTB1* (Os03g0706500), *OsT20* (Os12g0405200), *OsD27* (Os11g0587000), *OsD17* (Os04g0550600), *OsD10* (Os01g0746400), *OsD3* (Os06g0154200), *OsD14* (Os03g0203200), *OsPIN2* (Os06g0660200), *OsWOX11* (Os07g0684900), *OsRR1* (Os04g0442300), *OsRR2* (Os02g0557800), *OsRR3* (Os02g0830200), *OsRR4* (Os01g0952500), *OsRR5* (Os04g0524300), *OsRR6* (Os04g0673300), *OsRR7* (Os07g0449700), *OsRR9* (Os11g0143300), *OsRR10* (Os12g0139400), and *OsRR11* (Os02g0631700).

## RESULTS

### *OsCKX* family genes display different expression patterns

Similar to the findings of previous studies (Ashikari *et al*., 2005; Zhang *et al*., 2021), the *OsCKX* phylogenetic tree consisted of five major clades (Figure 1A). *OsCKX1*, which is commonly expressed in the roots, flowers, and grains, and *OsCKX2* were grouped together in the first clade. The β­glucuronidase (GUS) staining results revealed that *OsCKX1* had obvious expression in the shoot base and top of axillary buds (Figure 1B). On the other hand, *OsCKX2*, usually expressed at high levels in the leaf sheath and inflorescence, was found to be particularly expressed in the lateral root primordium, leaf blade, shoot base, and inflorescence. *OsCKX6*, *OsCKX7*, and *OsCKX10* were grouped together in the second clade and showed very low expression in all the tissues analyzed.

**Figure 1.**
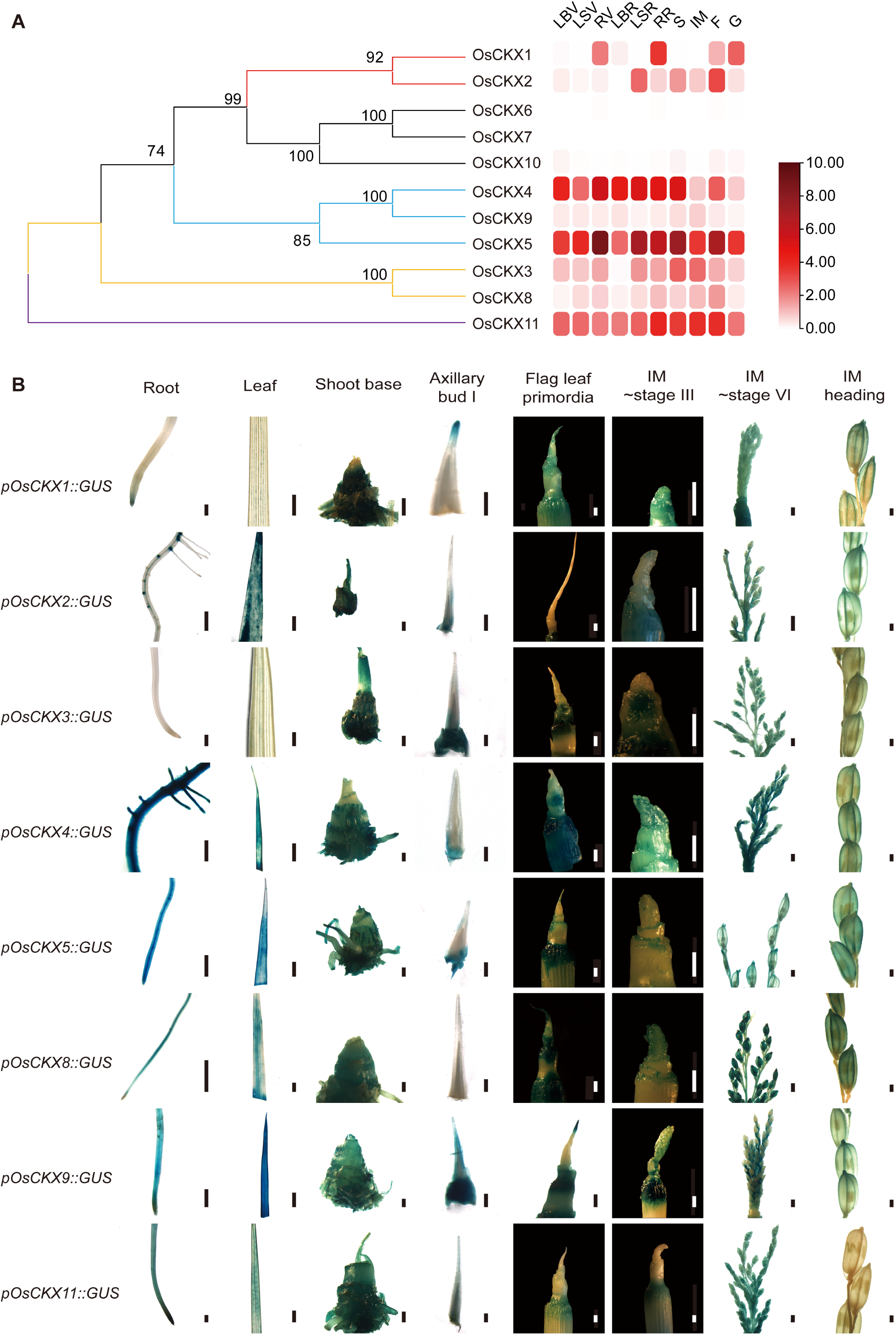
Genetic relationship and expression patterns of *OsCKX* genes based on RNA sequencing data and GUS staining assay. (A) Genetic relationship and expression patterns of *OsCKX* genes investigated in the leaf blade at the vegetative stage (LBV), leaf sheath at the vegetative stage (LSV), root at the vegetative stage (RV), leaf blade at the reproductive stage (LBR), leaf sheath at the reproductive stage (LSR), root at the reproductive stage (RR), stem (S), inflorescence meristem (IM), flower (F), and grain (G). The expression pattern was based on log2 (Fragments Per Kilobase of exon model per Million mapped fragments (FPKM) +1) values. (B) Histochemical GUS staining of *pOsCKX::GUS* transgenic plants at various developmental stages. The scale bar in B was 1 mm.

The third clade was composed of *OsCKX4*, *OsCKX5*, and *OsCKX9*. *OsCKX4*, which is generally strongly expressed in the vegetative organs, displayed extremely special expression in the roots. *OsCKX9* was generally expressed at low levels in all the tissues analyzed but showed special expression in the leaf blade and whole axillary buds. *OsCKX5*, usually highly expressed in all the checked tissues, also exhibited especial expression in the roots and leaves.

The fourth clade consisted of *OsCKX3* and *OsCKX8*. *OsCKX3*, which is expected to have the highest expression in the stem and young panicles, was found to have particular expression in the shoot base and young panicle. *OsCKX8*, commonly expressed at lower levels than *OsCKX3* in all vegetative organs, was extraordinarily expressed in the shoot base, flag leaf primordia, and inflorescence.

### *OsCKX11* showed a different sequence from other *OsCKX* genes; it was generally expressed at a higher level in all tissues, specifically in the reproductive-stage root and inflorescence, displaying special expression in the root, shoot base, and young inflorescence

Cytokinins have many functions during plant growth and development in different organs, and the biodegradation of cytokinins in different periods or tissues can help plants accurately regulate their growth and development, so that each *OsCKX* gene is expressed in a distinct pattern. In rice, we found all of eight of the *OsCKX* genes expressed at the shoot base and inflorescence meristem at different stages and positions. We found some tissues with low expression of certain *OsCKX* genes, such as *OsCKX2*, *OsCKX3*, and *OsCKX11*, which were not expressed in the flag leaf primordia. Additionally, we determined the expression patterns of *OsCKX* genes in axillary buds: *OsCKX1* was expressed at the top; *OsCKX2*, *OsCKX3*, *OsCKX4,* and *OsCKX5* were expressed at the base; and *OsCKX9* was expressed throughout the buds.

### *OsCKX* mutants exhibit different phenotypes

Previous studies suggested that the knockdown of specific *OsCKXs* might help improve rice production by influencing important agronomic traits, such as tillering, development of organs facilitating nutrition, and panicle phenotype. To confirm this hypothesis, we created mutants of all *OsCKX* genes using the CRISPR/Cas9 system, including single mutants of the nine *OsCKX* genes (except *OsCKX6* and *OsCKX10*); double mutants of *OsCKX1* and *OsCKX2* as well as *OsCKX4* and *OsCKX9*; and triple mutants of *OsCKX6*, *OsCKX7* and *OsCKX10* (Supplemental Figures S1-S3).

Since leaf size affects the photosynthetic capacity of rice, we investigated this and discovered that the top three leaves in the *osckx1*, *osckx2*, *osckx8*, and *osckx11* mutants were distinctly longer and wider compared to those of ZH11 (Supplemental Figures S4-S6; Supplemental Table S2). The last three leaves of the *osckx9* mutant were also distinctly longer, yet no change in their width was observed compared to ZH11. The leaf length and width of the other mutants did not significantly change (Supplemental Figures S5, A–D, and S6). On the other hand, culm diameter and plant height are closely related to the lodging resistance of rice. We found that the diameters of the basal internodes in *osckx1*, *osckx2*, *osckx8*, and *osckx11* mutants were significantly wider than those in the WT, while the *osckx3*, *osckx4*, and *osckx5* mutants had thinner basal internodes (Supplemental Figure S5, E and F). Furthermore, the *osckx3*, *osckx4*, and *osckx5* mutants showed significantly decreased plant height compared to others (Supplemental Figure S5, G and H).

The panicle number is determined by the tiller number per plant and has a significant effect on rice yield. In the field and pot experiments, the *osckx4* and *osckx9* mutants developed more tillers than the WT plants, which is consistent with the results of previous studies (Supplemental Figures S4B and S5, I and J; Supplemental Tables S2 and S3). Although the mutants produced more panicles per plant, the number of grains per panicle decreased, resulting in no significant change in yield (Supplemental Figure S7, A-D; Supplemental Table S2). In contrast, the *osckx2* mutant showed significantly reduced tiller number in both the 2019 and 2020 field and pot experiments (Supplemental Tables S2 and S3). However, no significant differences were observed in the tiller numbers of the other *osckx* mutants. Additionally, there was no significant change in the yield per plant of the *osckx* mutants, except for *osckx1* and *osckx11-1*, which had significantly improved production. In contrast, *osckx3-1*, *osckx3-26*, and *osckx4-6* had significantly low yields (Supplemental Figures S7, A and B). Despite this, we observed several agronomic traits that were improved in the mutants. For example, the *osckx1*, *osckx2*, *osckx7,* and *osckx8* mutants presented heavier grain weight, while the *osckx11* mutant developed more spikelets per panicle (Supplemental Figure S7, E–H).

Though OsCKX6, OsCKX7, and OsCKX10 have similar protein sequences, the three genes have very low expression levels. Using these genes, we constructed *osckx6 osckx7 osckx10* triple mutant lines; all the three lines showed no significant difference in panicle number and yield compared to the WT (Supplemental Figures S8).

### The *osckx1 osckx2* double mutants have significantly reduced tiller numbers

*OsCKX1* and *OsCKX2* displayed similar trends in the phenotypes of flag leaf length and width, culm diameter, and 1000-grain weight, and the two genes were grouped into the same clade. Based on this, we constructed the *osckx1 osckx2* double mutants. In 2021, both of the two *osckx1 osckx2* double mutant lines showed a significant increase in the number of spikelets per panicle in relation to NIP; this increase is mainly attributed to the significantly increased number of primary branches per panicle; however, the *osckx1 osckx2-19* mutant showed a significant increase in the number of secondary branches per panicle (Figure 2, A–D). A similar phenotype was also seen in 2020 (Supplemental Figure S9, A–D). However, compared to NIP, the seed setting rate of *osckx1 osckx2* mutant lines was significantly lower in the two years (Figure 2E, Supplemental Figure S9E). Furthermore, the panicle number of *osckx1 osckx2* mutant lines was less than that of NIP in 2021, while the *osckx1 osckx2-19* mutant showed a significantly reduced number of tillers and panicles per plant compared to NIP in 2020 (Figure 2F, Supplemental Figure S9, F and G). Moreover, the 1,000-grain weight and grain length were significantly increased in *osckx1 osckx2-19* compared with those in NIP (Supplemental Figure S9, H–J). In contrast, the grain width and thickness did not show significant changes between *osckx1 osckx2-19* and NIP (Supplemental Figure S9, K and L). Owing to the small panicle number per plant and lower setting rate, the yield of *osckx1 osckx2* lines was lower compared to that of NIP (Figure 2G, Supplemental Figure S9M). Moreover, in *osckx1 osckx2-19* plants, the flag leaf became wider (Supplemental Figure S9, N and O) and the plants had thick basal internodes, while the plant height did not noticeably change (Supplemental Figure S9, P and Q). Collectively, these observations of the spikelet phenotype, panicle number, and vegetative organ size suggest that the double knockout of *OsCKX1* and *OsCKX2* may help improve the lodging resistance and optimize the plant architecture in rice.

**Figure 2.**
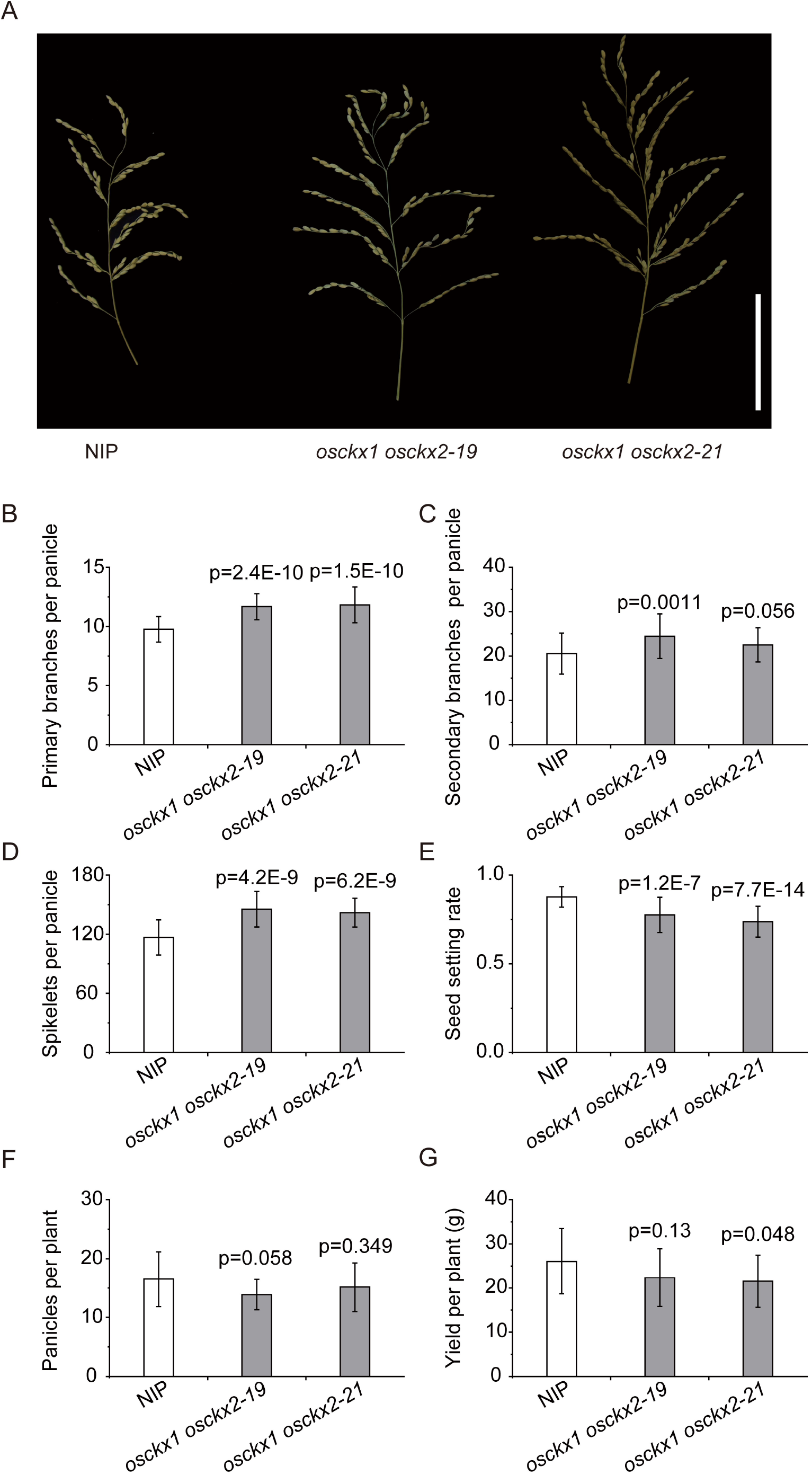
Phenotypic characterization of Nipponbare (NIP) and osckx1 osckx2 mutant plants in 2021. (A) Panicle phenotypes of NIP and *osckx1 osckx2* mutants. The image was digitally extracted and scaled for comparison (scale bar = 10 cm). (B–G) Measurement of the primary branches per panicle (B), secondary branches per panicle (C), spikelets per panicle (D), seed setting rate (E), panicles per plant (F) and yield per plant (G) of NIP and *osckx1 osckx2* mutants (n > 30 for B–E, n > 15 for F and G). P values indicate the level of statistical significance between NIP and *osckx1 osckx2* mutants determined using the Student’s *t*-test.

### Double knockout of *OsCKX4* and *OsCKX9* significantly promotes tillering

For *osckx4*, we observed four independent lines of the NIP background, and for *osckx9*, we observed two independent lines of the ZH11 background; all of these lines showed a relative phenotype in increased tiller numbers (Supplemental Figures S5, I and J; Supplemental Tables S2 and S3) and reduced spikelets (Supplemental Figure S7, E and F; Supplemental Table S2) per panicle. Moreover, OsCKX4 and OsCKX9 were grouped into the same clade. Based on these results, we mutated *OsCKX9* in NIP and *osckx4-6* mutants. The double mutant *osckx4 osckx9* has the same mutations as *osckx4-6* and *osckx9-1* (Supplemental Figures S3B). At the T0 generation, we obtained 5 lines of an *osckx4 osckx9* double mutant. All mutant lines showed a higher number of tillers than the wild type. We used two independent lines with the same mutations for further research. As expected, we discovered that the *osckx4 osckx9* double mutant had an extremely higher tiller number compared to NIP and the two single mutants (Figure 3, A–D). In contrast, panicle size was significantly reduced in the *osckx4 osckx9* mutant (Figure 3E). The *osckx9-1* mutant had a high panicle number per plant, which is the same as the *osckx9* single mutants in ZH11 background, but the number was markedly higher in the *osckx4 osckx9* mutant (Figure 3F). However, the number of spikelets per panicle of *osckx4* was significantly decreased, while that of *osckx4 osckx9* was even lower (Figure 3G). The decreased number of spikelets per panicle of *osckx4-6* was due to the low number of secondary branches per panicle, while the low number of spikelets per panicle of *osckx4 osckx9* was caused by the reduced number of both the primary and secondary branches (Figure 3, E, H, and I). On the other hand, seed setting rates were similar in the single and double mutants, and were significantly lower than those in WT (Figure 3J). Despite the increase in grain length, the *osckx4-6* mutant presented decreased 1,000-grain weight due to reduced grain thickness (Figure 3K; Supplemental Figure S10, A–E). The 1,000-grain weight of *osckx4 osckx9* was lower than that of the *osckx4-6* mutant due to reduced grain width and thickness (Figure 3K; Supplemental Figure S10, A-E). As a result of these changes, the yield of *osckx4 osckx9* was significantly decreased (Figure 3L). Moreover, the root length, diameter of basal internode, plant height, and flag leaf length and width were further decreased in the double mutant compared to those in the two single mutants (Figure 3, A, M and N; Supplemental Figure S10, F-H).

**Figure 3.**
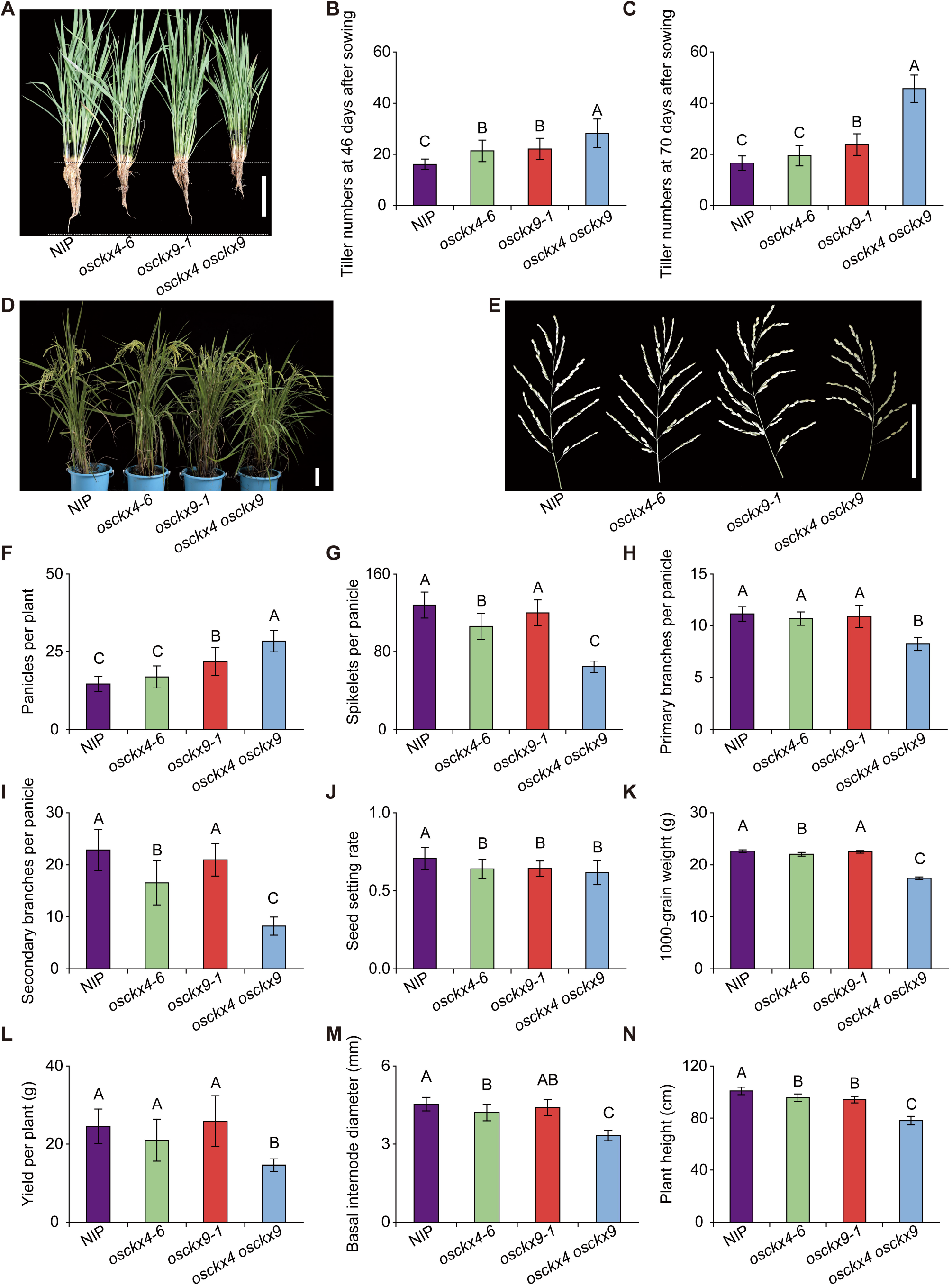
Phenotypic characterization of Nipponbare (NIP) and *osckx4*, *osckx9*, and *osckx4 osckx9* mutant plants from the field experiment. (A) Phenotypic features of NIP, *osckx4*, *osckx9* and *osckx4 osckx9* seedlings at the vegetative stage 46 days after sowing. The image was digitally extracted and scaled for comparison (scale bar = 10 cm). (B, C) Number of tillers at the vegetative stage 46 days after sowing (B) and reproductive stage 70 days after sowing (C) (n > 20 for NIP, *osckx4-6*, and *osckx9-1*, n = 10 for *osckx4 osckx9*). (D, E) Habits (D) and panicle phenotypes (E) of NIP, *osckx4-6*, *osckx9-1*, and *osckx4 osckx9* plants at the mature stage. The image was digitally extracted and scaled for comparison (scale bar = 10 cm). (F–N) Measurement of the panicle number (F), spikelets per panicle (G), primary branches per panicle (H), secondary branches per panicle (I), seed setting rate (J), 1,000-grain weight (K), yield per plant (L), diameter of basal internode (M), and plant height (N) of NIP, *osckx4-6*, *osckx9-1* (n > 20 for F–H and J–N, n = 5 for I), and *osckx4 osckx9* (n > 10 for F–J and L–N, n = 5 for K). Different capital letters represent the level of statistical significance (*P* < 0.01) determined using one-way ANOVA and shortest significant ranges (SSR) test.

We also conducted a hydroponic experiment to investigate the root phenotypes of the mutants (Figure 4A). We verified that *OsCKX4* and *OsCKX9* have redundant roles in regulating tillering, plant height, and root length (Figure 4 B and C). The *osckx4 osckx9* mutant not only had shorter roots, but also fewer crown roots and smaller root diameter (Supplemental Figure S11, A and B). The length and width of the 8^th^ leaf were also decreased in *osckx4-6* and *osckx4 osckx9* compared to those in NIP (Figure 4, D and E). These results indicate that the reduction in the leaf size of *osckx4 osckx9* started at the vegetative stage. Overall, these results suggest that *OsCKX4* and *OsCKX9* have a functional overlap during rice growth and development.

### RNA-seq analysis reveals genes with altered expression of *osckx4*, *osckx9*, and *osckx4 osckx9*

To obtain information regarding the possible functional context of *OsCKX4* and *OsCKX9*, we sampled the roots and shoot bases from NIP, *osckx4*, *osckx9*, and *osckx4 osckx9* and performed RNA-seq analysis. Results revealed that *osckx4* had 297 differentially expressed genes (DEGs) in the shoot bases and 415 DEGs in the roots compared to NIP. There were 500 DEGs in the shoot bases and 342 DEGs in the roots of the *osckx9* mutant compared to NIP. The *osckx4 osckx9* double mutant had the highest number of DEGs, with 806 DEGs in the shoot base and 4,556 DEGs in the roots compared to NIP (Figure 5A). To verify the accuracy and reproducibility of the RNA-seq results, we randomly selected seven previously studied genes associated with the phenotypes observed in this study for quantitative reverse-transcription PCR analysis. The expression profiles of these genes were found to corroborate the RNA-seq results (Supplemental Figure S12).

**Figure 4.**
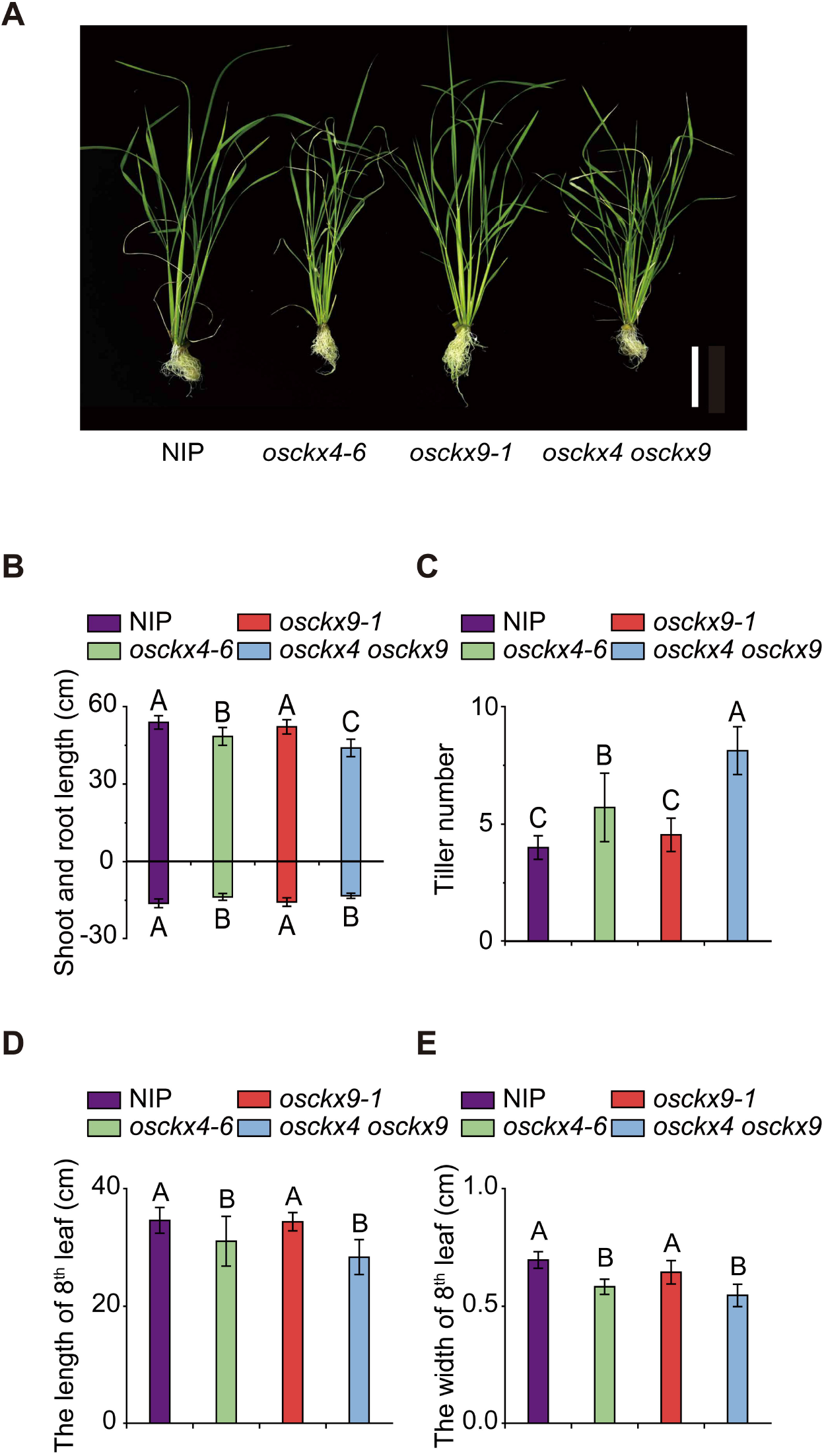
Phenotypic characterization of Nipponbare (NIP) and *osckx4*, *osckx9*, and *osckx4 osckx9* mutant plants from the hydroponic experiment. (A) Phenotypic features of Nipponbare (NIP), *osckx4*, *osckx9*, and *osckx4 osckx9* seedlings 21 days after transferring to hydroponic solutions. The image was digitally extracted and scaled for comparison (scale bar = 10 cm). (B–E) Measurement of the shoot (above X axis) and root (below X axis) lengths (B), tiller numbers (C), length of 8^th^ leaf (D), and width of 8^th^ leaf (E) of NIP, *osckx4-6*, *osckx9-1*, and *osckx4 osckx9* seedlings at 21 days after transplant (n = 24 each). Different capital letters represent the level of statistical significance (*P* < 0.01) determined using one-way ANOVA and SSR test.

**Figure 5.**
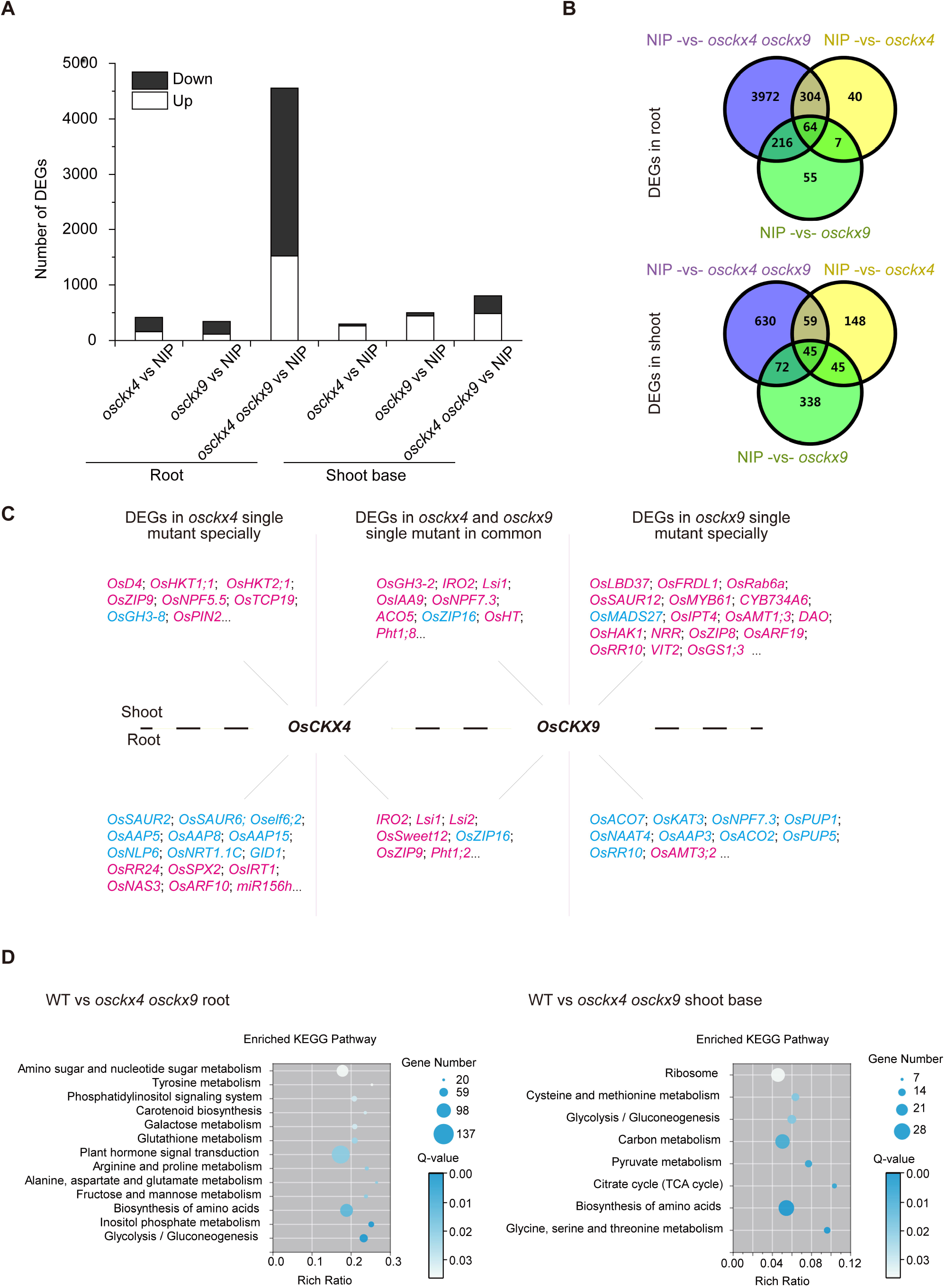
RNA-seq analysis of the roots and shoot bases (BP) of Nipponbare (NIP) and *osckx4*, *osckx9*, and *osckx4 osckx9* mutant plants. (A) The number of differentially expressed genes (DEGs) identified in the roots and shoot bases of the following pairwise comparisons: Nipponbare (NIP) vs. *osckx4*, NIP vs. *osckx9*, and NIP vs. *osckx4 osckx9*. The changes in gene expression levels were calculated using the log2 fold change and *Q* values from three biological replicates. (B) Overlapping DEGs in the roots and shoot bases (BP) of NIP vs. *osckx4*, NIP vs. *osckx9*, and NIP vs. *osckx4 osckx9*. (C) DEGs in *osckx4* and *osckx9* specifically and in common in the roots and shoots, genes in magenta are upregulated; genes in sky blue are downregulated. (D) Top enriched Kyoto Encyclopedia of Genes and Genomes (KEGG) pathways of the DEGs identified in the roots and shoot bases of NIP vs. *osckx4 osckx9*.

However, although there were numerous DEGs between the WT and mutants, there were 71 DEGs in the roots and 90 DEGs in the shoot bases that overlapped between the *osckx4* vs. WT and *osckx9* vs. WT comparisons and occupied only a small fraction of the total DEGs (Figure 5B). These included some genes that regulate the absorption of silicon (*Lsi1*, *Lsi2*), zinc (*OsZIP16*), and iron (*OsIRO2*) (Fig 5C). *OsZIP9*, *OsTCP19*, *OsHKT1.1*, and *OsNPF5.5* were upregulated in the shoots of *osckx4*, while *OsLBD37*, *OsARF19*, *OsMYB61*, and *MT1a* were upregulated in the shoots of *osckx9*. In the roots of *osckx4*, the expression of *OsAAP5*, *OsAAP8*, and *OsNLP6* was downregulated, while certain genes influencing iron and phosphorus absorption were upregulated. We also observed the downregulation of *OsPUP1*, *OsPUP5*, *OsRR10*, and *OsNAAT4* in the roots of *osckx9* (Figure 5C). However, the functions of most DEGs remain unknown (Supplemental Dataset 1). The difference in DEGs between the two single mutants and NIP implied that *OsCKX4* and *OsCKX9* have different functions at the molecular level. Moreover, 87.1% and 78.2% of DEGs in the roots and shoot bases, respectively, between the *osckx4 osckx9* and WT were identified only in the double mutants, which may have been caused by the duplicate effect of the functional loss of *OsCKX4* and *OsCKX9* (Figure 5B).

Since we observed that *osckx4 osckx9* exhibited more severe phenotypes and possessed more DEGs, we analyzed the DEGs identified in the roots and shoot bases between the WT and *osckx4 osckx9* plants by using Kyoto Encyclopedia of Genes and Genomes (KEGG) enrichment analysis (Figure 5D). In both tissues, the top enriched KEGG pathways included ‘glycolysis/gluconeogenesis’, ‘biosynthesis of amino acids’, and those related to carbon and nitrogen metabolism. Since most of the DEGs were not functionally annotated, the relationship between the DEGs and the enriched pathways and their influence on the growth and development of the mutant plants were not fully understood. However, we identified several genes that may be associated with the phenotypes observed in this study.

### OsCKX4 and OsCKX9 regulate cytokinin levels

As OsCKX4 and OsCKX9 are involved in biodegradation of cytokinins, we first analyzed the DEGs associated with cytokinin metabolism and signal transduction. Among the cytokinin metabolism-related genes, only *OsCKX2* and *OsCKX5* expression was downregulated in the root of *osckx4 osckx9* (Supplemental Dataset 1).

Type-A *OsRR* genes are a group of genes that can be induced by cytokinins. However, the expression of *OsRR4*, *OsRR6*, *OsRR7*, *OsRR9*, and *OsRR10* was significantly downregulated in the roots of *osckx4 osckx9*, whereas the expression of *OsRR10* was significantly downregulated in the roots of *osckx9* (Figure 6A). In the shoot base, only *OsRR10* expression was significantly upregulated in *osckx9* and *osckx4 osckx9* mutants (Figure 6A). In contrast, in the axillary bud, the expression of detectable type-A *OsRR* genes, except *OsRR2*, was significantly upregulated in *osckx4 osckx9* mutants (Figure 6B).

**Figure 6.**
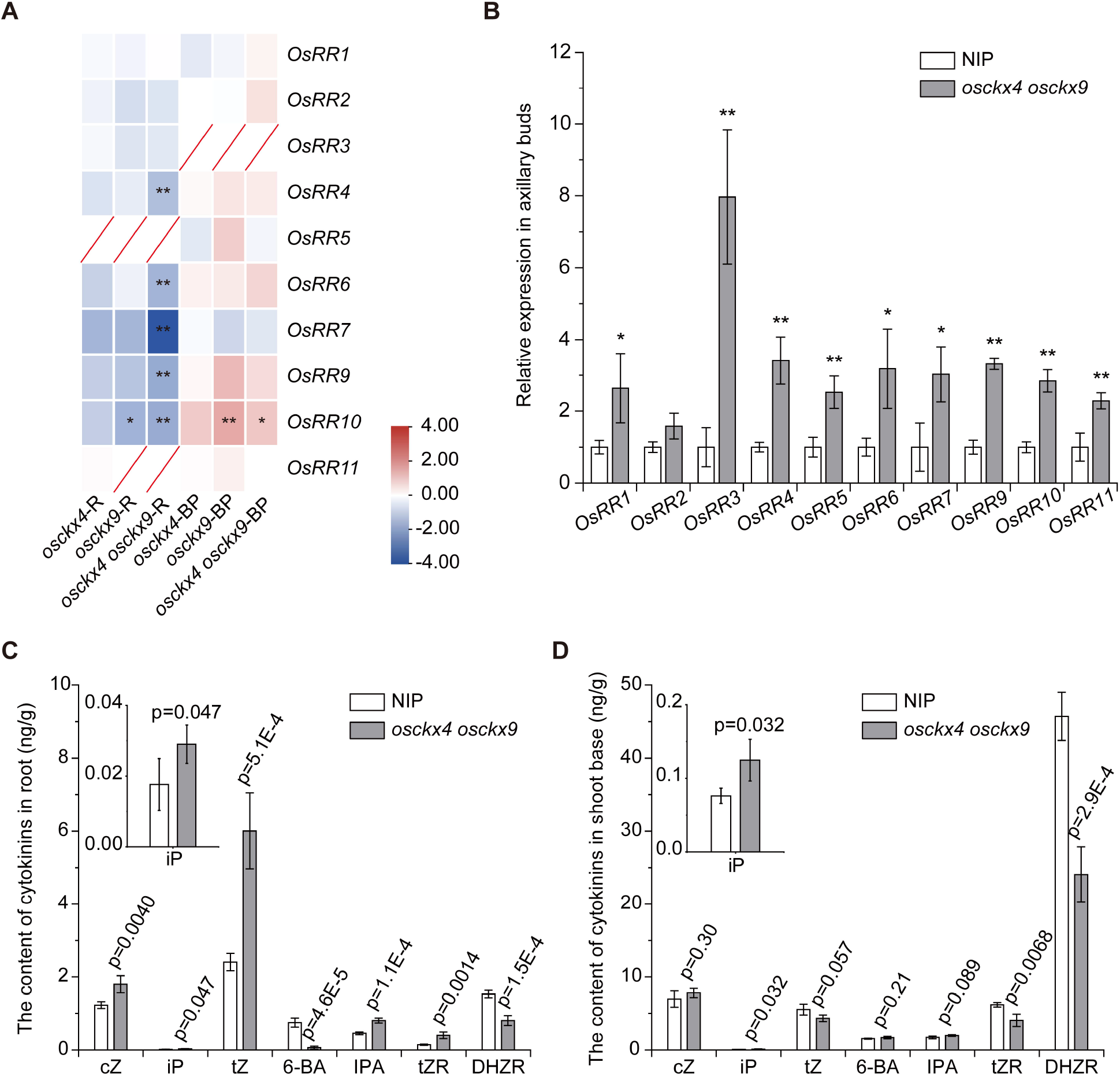
OsCKX4 and OsCKX9 can regulate endogenous cytokinin levels in plant tissues. (A) Differentially expressed type-A *OsRR* genes in the root (R) and shoot base (BP) of different mutants compared with Nipponbare (NIP). The fold change was based on log2 (FPKMmutant/FPKMNIP) values. * Q < 0.05, ** Q < 0.01. (B) Transcript levels of *OsRR* genes at 0–1 cm axillary buds determined using qRT-PCR. β-Actin was used as a reference gene. Results are presented relative to NIP. Data represent means ± SD (n = 3). P values represent the level of statistical significance between NIP and *osckx4 osckx9*. * P < 0.05, ** P < 0.01, determined using the student’s *t*-test. (C, D) The content of endogenous cytokinins in root (C) and shoot base (D) of 4-week­old seedlings. cZ (*cis*-zeatin), iP (*N*^6^-(Δ ^2^-isopentenyl) adenine), tZ (*trans*-zeatin), 6-BA (6-benzylaminopurine), IPA (isopentenyl adenosine), tZR (*trans*-zeatin riboside) and DHZR (dihydrozeatin riboside) were detected in this study. Data represent means ± SD (n = 4). P values indicate the level of statistical significance between NIP and *osckx4 osckx9* mutants determined using the Student’s *t*-test.

Although type-A *OsRR* genes can indicate the content of cytokinins indirectly, we sampled the roots and shoot base of NIP and *osckx4 osckx9* to detect the content of cytokinins. In the root, compared to NIP, the content of cZ (*cis*-zeatin), iP, tZ, IPA (isopentenyl adenosine), and tZR (*trans*-zeatin riboside) showed a significant increase, while 6-BA and DHZR (dihydrozeatin riboside) contents significantly decreased in *osckx4 osckx9* (Figure 6C). However, in the shoot base, we found a higher content of iP and lower content of tZR and DHZR in *osckx4 osckx9* (Figure 6D).

### SL metabolism and signaling are not regulated by *OsCKX4* and *OsCKX9*

SLs perform the function of regulating tiller numbers (Jiang *et al*., 2013; Zhou *et al*., 2013). To investigate the cause of the increased tiller numbers in the single and double mutants, we examined the genes associated with SLs. In the root and shoot base, none of the genes related to SL biosynthesis and signaling transduction, including *OsT20* (Liu *et al*., 2020), *OsD27*, *OsD17*, *OsD10*, *OsD3*, *OsD14* (Arite et al., 2007), and *OsD53* (Jiang *et al*., 2013; Zhou *et al*., 2013), showed no significant difference between NIP and mutants (Supplemental Figure S13; Supplemental Dataset 1). Based on these results, there is no direct evidence that the metabolism and signaling transduction of SLs are downstream of *OsCKX4* and *OsCKX9*.

### *OsCKX4* and *OsCKX9* may participate in regulating the metabolism and transport of other hormones

To investigate the cause for the increases in tillers and the reduced root length in *osckx4 osckx9*, considering the importance of auxin in regulating root elongation (Jun *et al*., 2011) and tiller bud outgrowth (Zha *et al*., 2019), we analyzed the genes related to auxin. We discovered that auxin efflux transporters *OsPIN2* was upregulated in the shoot bases, while *OsPIN1d* and *OsPIN8* were upregulated in the roots, respectively, in double mutants (Figure 7; Supplemental Dataset 1). We also found that the auxin primary response genes *OsGH3-2* and *OsIAA9* were upregulated in the shoot bases of the *osckx4 osckx9* and single mutants. Additionally, *OsGH3-2*, nine *OsIAA* genes (*OsIAA1, 4, 9, 12, 14, 15, 19, 23, 24*), and five *OsSAUR* genes (*OsSAUR6, 20, 27, 28, 33*) were downregulated in the roots of *osckx4 osckx9*; however, *OsIAA16*, *OsARF10* and *OsARF24* were upregulated in the roots of *osckx4 osckx9*. These results may partly explain the reduced root system and increased tiller numbers of *osckx4 osckx9* double mutants, but these conclusions are only based on RNA-Seq, more research is required to learn the relationship between cytokinins and auxins.

**Figure 7.**
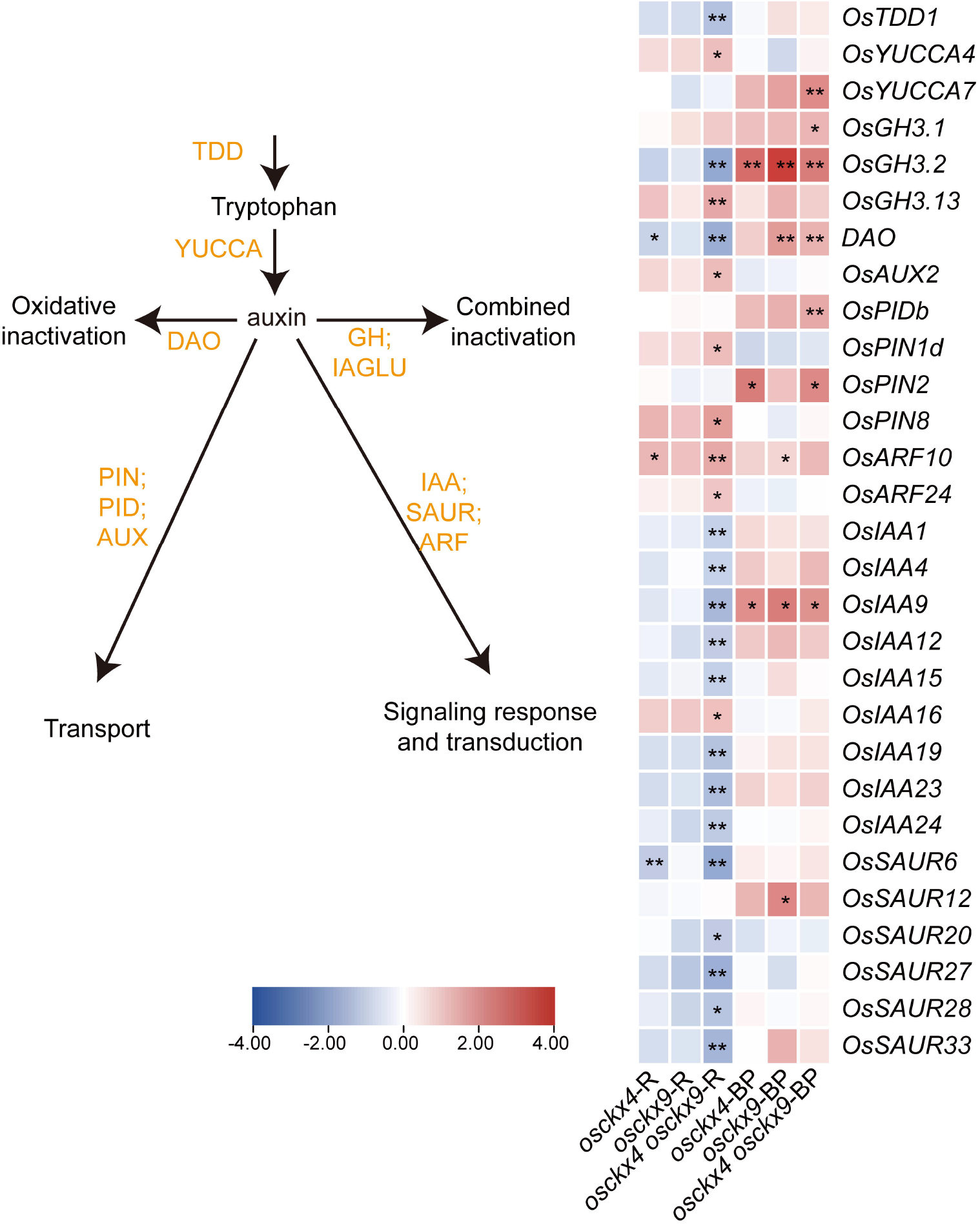
OsCKX4 and OsCKX9 are associated with auxin metabolism and signaling transduction. Differentially expressed genes (DEGs) related to auxin metabolism and signaling transduction in the root (R) and shoot base (BP) of different mutants compared with Nipponbare (NIP). The fold change was based on log 2 (FPKMmutant/FPKMWT) values. * *Q* < 0.05, ** *Q* < 0.01.

Additionally, to explain the reason for reduced plant height of *osckx4 osckx9*, we identified genes related to gibberellin metabolism, as gibberellins can regulate plant height (Sasaki *et al*., 2002). We found that the expression of *OsKS1* and *OsKS2* was significantly downregulated in the roots of *osckx4 osckx9* (Figure 8; Supplemental Dataset 1). OsKS1 and OsKS2 are enzymes that catalyze the second step of the GA biosynthesis pathway (Ji *et al*., 2014; Tezuka *et al*., 2015). The expression of *OsKS2* was also downregulated in the roots of the *osckx9* mutant, while no changes were observed in the *osckx4* mutant. Other genes for GA biosynthesis and degradation, such as *GA20ox* and *GA2ox*, were markedly downregulated in the roots of the *osckx4 osckx9*. Some GA receptors were also downregulated in the roots of the *osckx4 osckx9*. Although the changes in the expression of gibberellin­related genes may explain the significantly reduced plant height of *osckx4 osckx9* double mutants, deeper explore is needed to explain the crosstalk between cytokinins and gibberellin.

**Figure 8.**
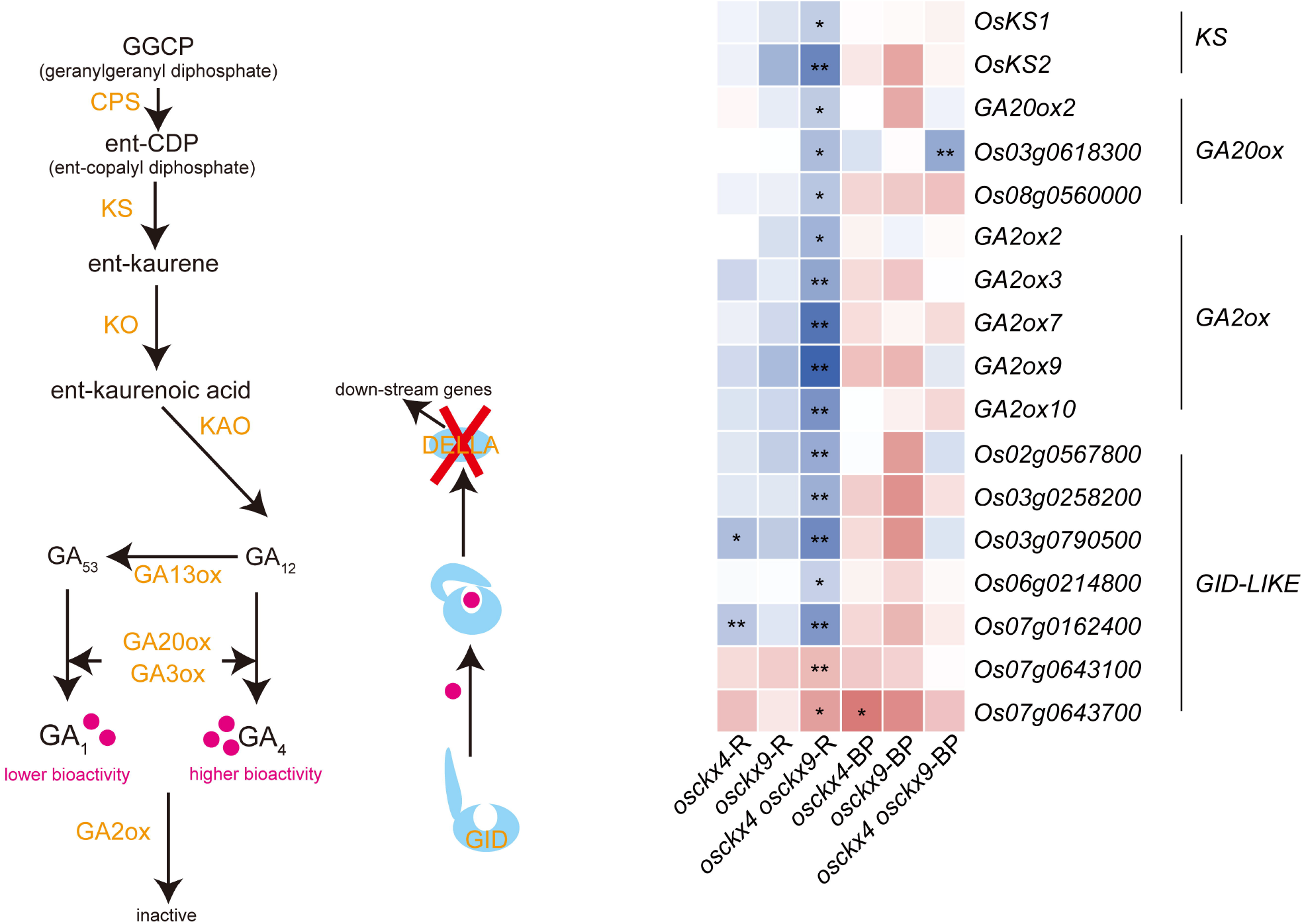
OsCKX4 and OsCKX9 are associated with gibberellin metabolism and signaling transduction. Differentially expressed genes (DEGs) related to gibberellin metabolism and signaling transduction in the root (R) and shoot base (BP) of different mutants compared with Nipponbare (NIP). The fold change was based on log2 (FPKMmutant/FPKMWT) values. * *Q* < 0.05, ** *Q* < 0.01.

## DISCUSSION

### Functional differences and redundancy between *OsCKX* genes

To systematically determine the functions of *OsCKX* genes, we analyzed their expression patterns and investigated the phenotypes of *osckx* single, double and triple mutation lines. Our study revealed the roles of all *OsCKX* genes in the growth and development of rice. Each *OsCKX* gene had its own special functions (summarized in Figure 9). Loss of function of some *OsCKX* genes cause changes to the yield.

**Figure 9.**
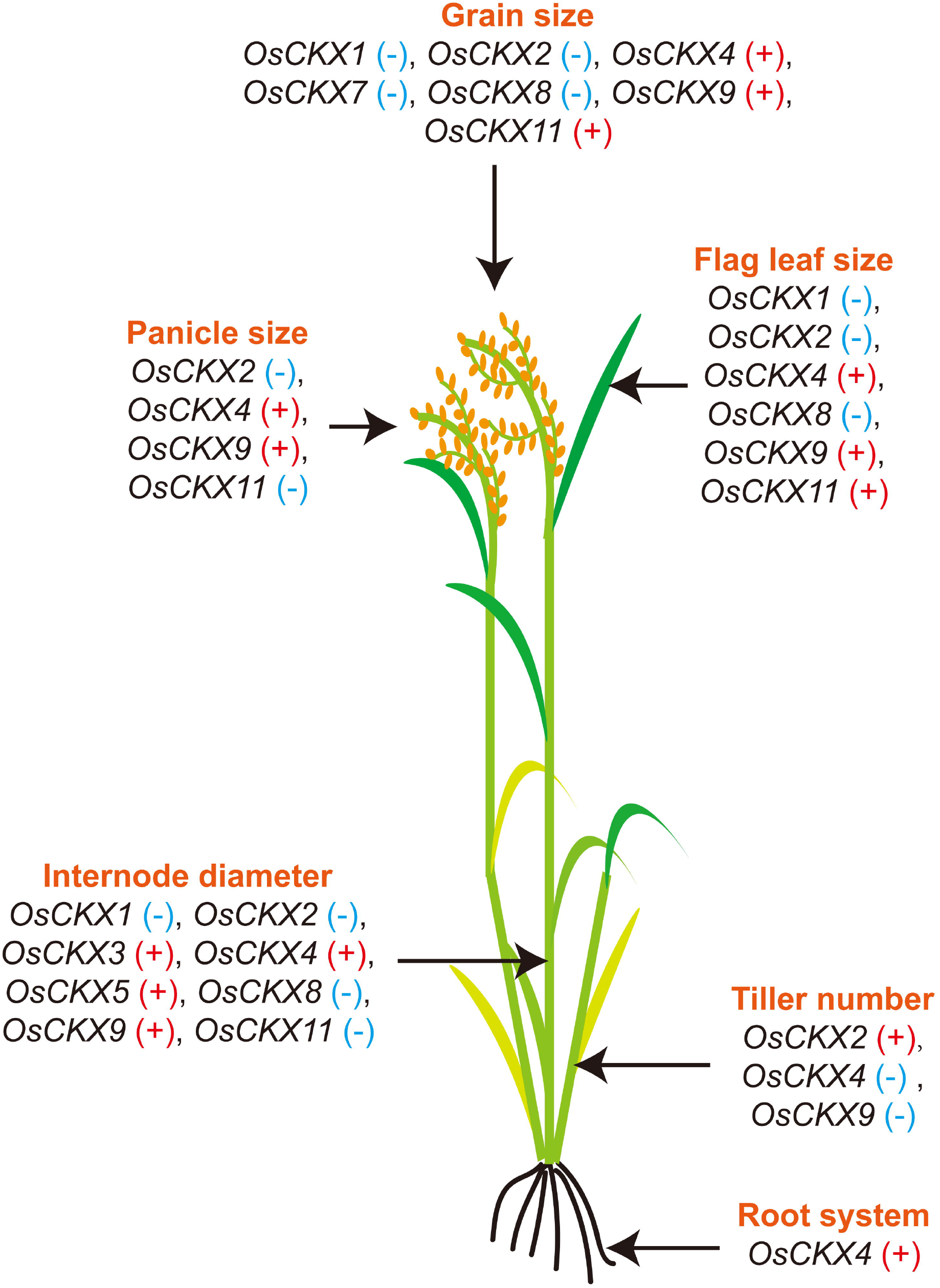
Functions of *OsCKX* genes in rice. The function of each *OsCKX* gene in regulating the growth and development of each organ in rice summarized from the phenotypes of all mutants. “+” means positive regulation, while “-” means negative regulation.

Previous studies have described the functions of *OsCKX2*, *OsCKX4*, *OsCKX9*, and *OsCKX11*. First, researchers elucidated the function of *OsCKX2* after observing that null mutants or plants with downregulation of *OsCKX2* developed more spikelets per panicle (Ashikari *et al*., 2005), more tillers, and larger grains (Yeh *et al*., 2015). Our present study observed little increase in the number of spikelets, but reduced tiller number in the *osckx2* mutant of ZH11, which were different from previous studies. Similar results of tiller numbers were also found in *osckx1 osckx2* double mutant of NIP, which supports the hypothesis that fewer tillers may be related to thicker culms. Furthermore, both the *osckx2* mutant of ZH11 and *osckx1 osckx2* mutant of NIP presented increased 1,000-grain weight. The discrepancies between our results and previous findings regarding panicle size may be attributed to the different cultivated varieties or environmental impact.

Furthermore, previous reports stated that *OsCKX4* showed a positive impact on the root system but a negative impact on the shoot growth (Gao *et al*., 2014; Mao *et al*., 2020). Since *OsCKX4* is commonly expressed in the root, previous studies were more focused on investigating the relationships between *OsCKX4* and the root system instead of the shoot system. Here, we discovered that *osckx4* mutants produced more tillers, especially at the early stages, but has fewer spikelets per panicle. Additionally, *osckx9* mutants have been reported to develop more tillers, fewer spikelets per panicle, and shorter plant height (Duan *et al*., 2019). Since the null mutations of *OsCKX4* and *OsCKX9* can affect the tiller number, we constructed *osckx4 osckx9* mutants, which distinctly showed more tillers, shorter plant height, and smaller panicles. The increase in the tiller number of *osckx4 osckx9* double mutants was greater than observed in *osckx4* and *osckx9* single mutants. Furthermore, the decrease in plant height and number of spikelets per panicle of *osckx4 osckx9* double mutants was also higher than observed in *osckx4* and *osckx9* single mutants. These results suggest the partial functional overlap of *OsCKX4* and *OsCKX9* during rice growth and development. Moreover, *osckx11* mutants previously displayed more tillers, larger panicles, and lighter 1,000-grain weight (Zhang *et al*., 2020); these were also observed in our study.

In contrast to our expectations, none of the single mutants showed a stable significant increase in yield, while the double mutants *osckx1 osckx2* and *osckx4 osckx9* showed lower yields. The lower yield is correlated with some of the disadvantageous phenotypes in these mutants. The discoveries in this study pointed out that single loss-of-function mutants were not good for breeding. If the expression of *OsCKX* gene in different tissues or at different developmental stages can be regulated in breeding, some favorable phenotypes may be utilized. Potential causes of the functional differences and redundancy in *OsCKX* genes Each *OsCKX* gene has its distinct functions during rice growth and development, but why different phenotypes arise is worthy of discussion. The main function of CKX enzymes is the biodegradation of cytokinins, with previous studies reporting that CKXs catalyze the degradation of different cytokinin compounds. In maize, the ZmCKX enzyme activity has been systematically analyzed; for example, ZmCKX1 and ZmCKX12 specifically target iP, tZ, and cZ (Zalabak *et al*., 2014). In *Arabidopsis*, like AtCKX1 showed the highest preference for tZ phosphates, iP phosphates, and iP9G, while AtCKX3 preferred iP phosphates and iPR (Kowalska *et al*., 2010). However, little is known about the preference of OsCKX enzymes; an in vitro experiment showed that OsCKX9 had higher activity to catalyze the degradation of tZ than to catalyze the degradation of IPA, and OsCKX11 that specially targets tZ and cZ (Duan *et al*., 2019; Zhang *et al*., 2020). While other OsCKX enzyme substrates may need to be inferred from endogenous hormone content data. From the hormone content detected in the root of *OsCKX4*-overexpression line, it was found that the content of iP and tZ cytokinin showed no obvious change compared to WT, while DHZ and the combined type of iP and tZ showed a significant reduction (Gao *et al*., 2014). In our study, in the root and shoot base, not all kinds of cytokinins showed higher levels in *osckx4 osckx9* mutants. Therefore, the distinctive functions of OsCKXs may be attributed to their different target substrates.

CKX proteins are mainly localized in the ER because most contain an N-terminal signal peptide sequence (Niemann *et al*., 2018; Zalabak *et al*., 2016). In the ER, cytokinins combine with their receptors and initiate signaling transduction, while most ER-localized CKXs regulate cytokinin concentrations to precisely adjust cytokinin signals (Niemann *et al*., 2018; Romanov and Schmulling, 2021). Previous studies revealed that OsCKX4 and OsCKX11 are localized in the cytosol, while OsCKX9 is localized in the cytosol and nuclei (Duan *et al*., 2019; Gao *et al*., 2014; Zhang *et al*., 2020). These three CKXs potentially regulate the cytokinin balance in other cellular organelles. However, the locations of the remaining OsCKXs remain unknown. Hence, we hypothesize that the subcellular localization of OsCKXs may also influence their functional differentiation.

In this study, the *OsCKX* genes exhibited different expression patterns. For example, *OsCKX1* and *OsCKX2* were expressed in the flower and grains, but *OsCKX1* showed lower expressed in the inflorescence meristem, which is the same as the previous study (Ashikari *et al*., 2005). As *OsCKX4* and *OsCKX9* showed similar phenotypes in tiller number, the expression pattern showed a considerable difference. *OsCKX4* is highly expressed in vegetative organs, yet the expression of *OsCKX9* in these organs is very low, and the different expression can get illation from previous studies (Duan *et al*., 2019; Gao *et al*., 2014). In inflorescence meristem, the two genes showed low expression level, which is also the same as previous studies by RT-southern blot (Ashikari *et al*., 2005). In summary, the differences in the nature of catalytic substrates, subcellular localization, and expression in tissues may contribute to the different functions observed in *OsCKX* family genes.

### Significantly altered transcriptome of the *osckx4 osckx9* mutant

Although *OsCKX4* and *OsCKX9* have been extensively studied, we observed new phenotypes in the single mutants, which have not been reported previously. Interestingly, these two genes were found to exhibit redundant functions during rice growth and development.

First, we found that several type-A *OsRR* genes showed lower expression in the roots in the *osckx4 osckx9* mutant compared to NIP, while no significantly changes (except *OsRR10*) in the shoot base. However, considering the higher content of most types of cytokinins in the root of the *osckx4 osckx9* mutant, these results were contradictory, since type-A *RR* genes are primary response genes for cytokinin signaling. Such results were not observed in the shoot base. Interestingly, similar results were also reported in some previous studies; for example, compared to those in WT, a lower content of cytokinins was recorded in the roots of *OsCKX4*-overexpression lines, with higher expression of *OsRR4*, *OsRR9*, and *OsRR10* (Gao *et al*., 2014). On the other hand, compared to those in WT, a higher cytokinin content was recorded in the roots, and lower in the shoots of the mutant *osabcg18*, but the expression of *OsRR6* and *OsRR10* in the roots did not differ significantly; in addition, the expression of *OsRR2, OsRR6 and OsRR10* in the shoots were downregulated (Zhao *et al*., 2019). These results may be caused by the response mechanism divergence between endogenous and exogenous cytokinins, the different functions of each type of cytokinin, and the distinct phenotypes and response pathways of endogenous cytokinins between the root and shoot. However, these ideas need to be verified by further research. Finally, the upregulated expression of nine type-A *OsRR* genes in the axillary buds of *osckx4 osckx9* may have resulted from the increase in the levels of cytokinins; however, owing to the limitations of the technique used, the levels of cytokinins in this tissue remains unknown.

Though higher cytokinin levels in the *osckx4 osckx9* mutant can partly explain the increased tiller number, more information about other plant hormones may be helpful for the analysis. However, from the RNA-Seq analysis, we did not observe any changes in the expression of SL biosynthesis-and signaling-related genes in the shoot base or the root. However, the higher expression level of the *OsPIN2* gene in the shoot base of *osckx4* and *osckx4 osckx9* mutants may help produce more tillers (Chen *et al*., 2012). Since SL directly activates *OsCKX9* to regulate shoot architecture in rice, based on these findings, cytokinin may be located downstream of the SL pathway during rice tillering. Further, though previous studies revealed some relationships between cytokinin content (Gao *et al*., 2014) or cytokinin signaling (Simaskova *et al*., 2015; Waldie and Leyser, 2018) and the expression of *PIN* genes, the connection between cytokinins and auxin in regulating tiller numbers require further research.

The reduced plant height of the *osckx4 osckx9* mutant may be related to the downregulation of *OsKS1* and *OsKS2*. On the other hand, possibly due to the limited substrate, the expression of several *GA20ox* and *GA2ox* genes for gibberellin biosynthesis and biodegradation, respectively, were downregulated in the *osckx4 osckx9* mutant. However, the mechanism by which *OsCKX4* and *OsCKX9* regulate the expression of *OsKS1* and *OsKS2* is unknown.

Interestingly, in addition to the genes regulated by both *OsCKX4* and *OsCKX9*, we also identified several genes that are regulated by *OsCKX4* and *OsCKX9* individually. In summary, the similar amino acid sequences but different expression patterns and different subcellular localization of *OsCKX4* and *OsCKX9* enable them to functionally overlap in the regulation of some pathways, while also individually regulating specific pathways.

In conclusion, our systematic analysis of 11 genes in the *OsCKX* family reveals their complex expression patterns in rice. A total of 27 *OsCKX* mutant lines were produced using CRISPR/Cas9 technology. By examining the phenotypes of the mutants throughout the rice growth period, we determined the functions of specific *OsCKX* genes in plant development. Specifically, we discovered that *OsCKX4* and *OsCKX9* inhibited tillering, while *OsCKX1* and *OsCKX2* promoted tillering. Considered together, our findings establish a community resource for fully elucidating the function of *OsCKXs*, providing new insights that may be used for future studies to improve rice yield.

## Supplemental data

The following supplemental materials are available.

**Supplemental Figure S1.** Gene structures and mutation details of *OsCKXs*.

**Supplemental Figure S2.** Chromatograms of the *osckx* mutant lines.

**Supplemental Figure S3.** Mutation details and chromatograms of double and triple mutants

**Supplemental Figure S4.** The *osckx* mutants of Zhonghua11 from the 2020 field experiment.

**Supplemental Figure S5.** Phenotypic characterization of the vegetative organs in *osckx* mutants from the 2020 field experiment.

**Supplemental Figure S6.** Phenotypic characterization of the second and third top leaves in *osckx* mutants from the 2020 field experiment.

**Supplemental Figure S7.** Phenotypic characterization of yield-related phenotypes in *osckx* mutants from the 2020 field experiment.

**Supplemental Figure S8.** Phenotypic characterization of the yield and panicles per plant in Nipponbare (NIP) and *osckx6 osckx7 osckx10*plants.

**Supplemental Figure S9.** Phenotypic characterization of Nipponbare (NIP) and *osckx1 osckx2-19* mutant plants in 2020.

**Supplemental Figure S10.** Phenotypic characterization of the flag leaves and seeds in Nipponbare (NIP), *osckx4*, *osckx9*, and *osckx4 osckx9* plants.

**Supplemental Figure S11.** Phenotypic characterization of the leaf and root systems in wild-type and *osckx4 osckx9* plants.

**Supplemental Figure S12.** Relative expression levels and FPKM values of *D17*, *RR4*, *WOX11*, *PIN2*, *TB1*, *OsMADS25*, and *OsMADS57* in the roots and shoot bases (BP) of Nipponbare (NIP), *osckx4*, osckx9, and osckx4 osckx9.

**Supplemental Figure S13.** OsCKX4 and OsCKX9 are associated with strigolactone biosynthesis and signaling transduction.

**Supplemental Table S1.** Information of primers used in the study.

**Supplemental Table S2.** Characterization of the vegetative organs and yield-related phenotypes in *osckx* mutants from the 2019 field experiment.

**Supplemental Table S3.** Tiller numbers in *osckx* mutants from the 2019 field experiment. Supplemental Dataset 1. RNA sequencing data.

## Data availability

All data supporting the findings of this study are available within the paper and within its supplementary data published online.

## Acknowledgments

We thank Jiankang Zhu and Caixia Gao for providing the vectors of the CRISPR-Cas9 system. We also thank Biogle and Biorun genome editing center for producing transgenic rice.

## Author contributions

C.D. and Y.D. conceived the original screening and research plans; C.R., Y.L., Z.C., Z.L., and C.D. performed the experiments and analyzed the data; C.R. and C.D. wrote the article with contributions of all the authors; C.D. agrees to serve as the author responsible for contact and ensures communication.

