## Supplemental figures and tables for "Cytokinin oxidase/dehydrogenase family genes exhibit functional divergence and overlap in rice growth and development, especially in control of tillering"

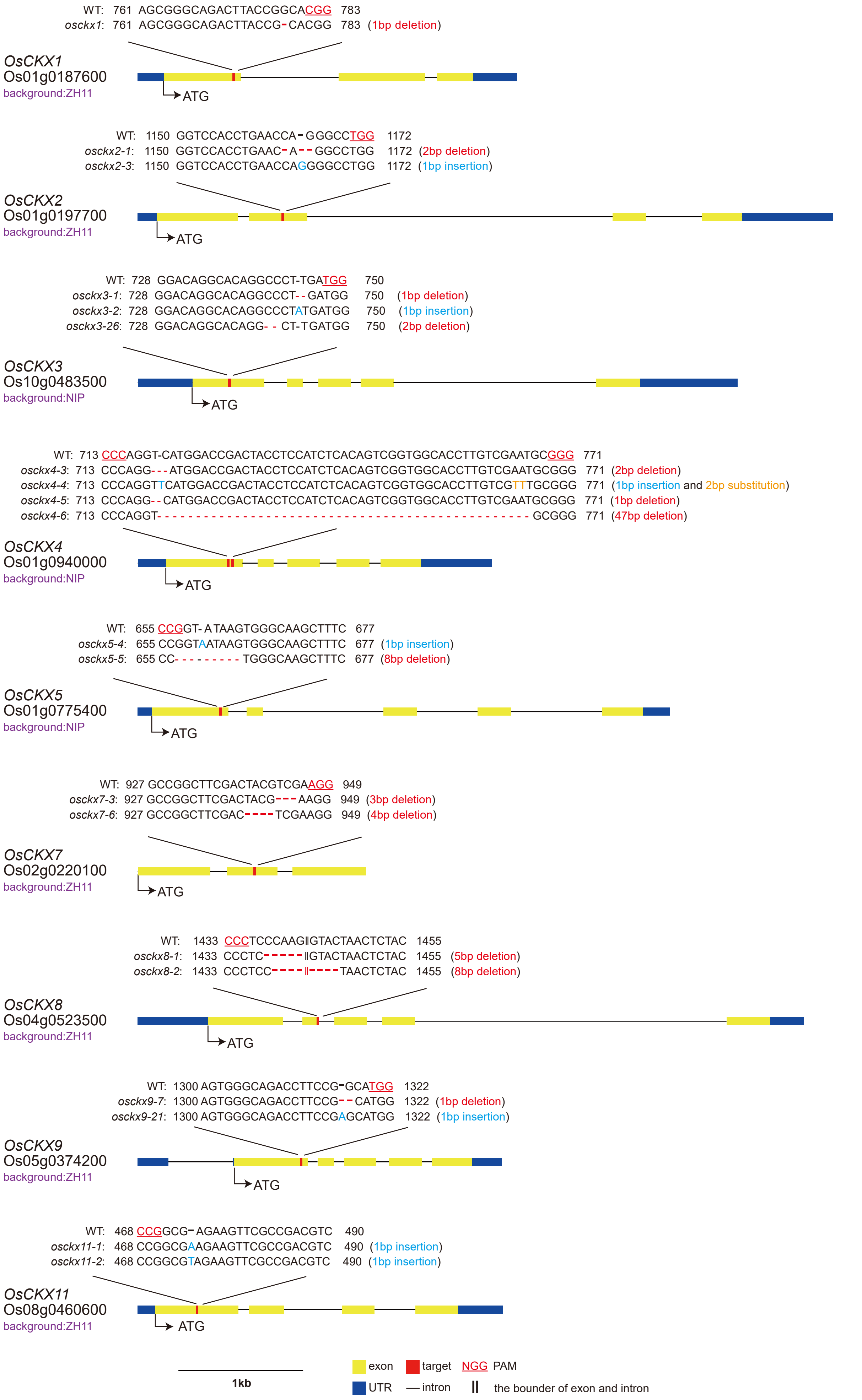


**Supplemental Figure S1.** Gene structures and mutation details of *OsCKX*s. Gene structures and mutation details of *OsCKX*s. The mutation details of the CRISPR edited lines targeting *OsCKX1*, *OsCKX2*, *OsCKX3*, *OsCKX4*, *OsCKX5*, *OsCKX7*, *OsCKX8*, *OsCKX9* and *OsCKX11*. Solid yellow boxes represent exons, solid lines represent introns, solid blue boxes represent untranslated regions, red boxes represent target sequences, and protospacer adjacent motif (PAM) sequences are shown in red with a solid red line underneath.

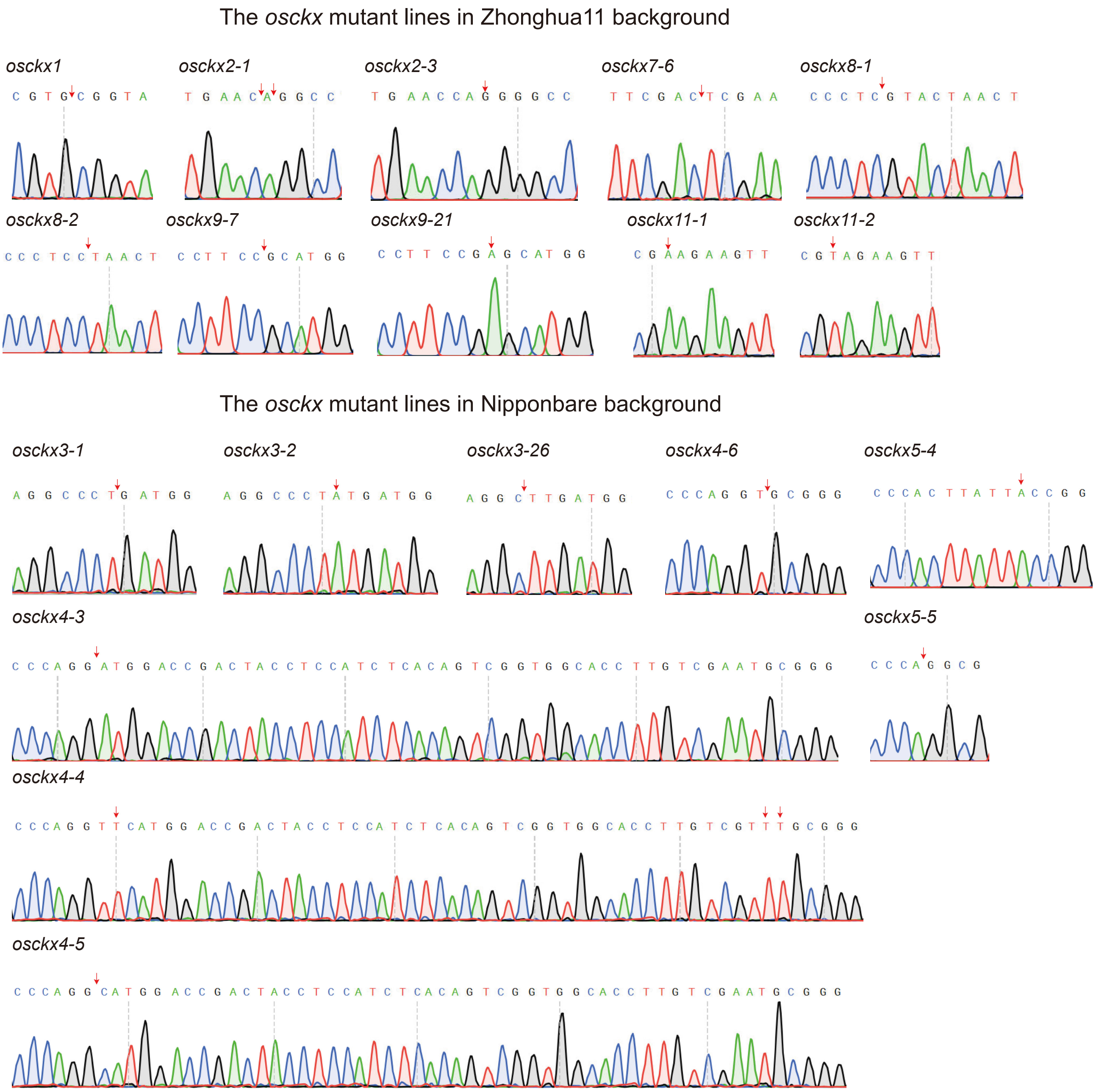


**Supplemental Figure S2.** Chromatograms of the *osckx* mutant lines. The sequencing results of each mutation line. Red arrows point to mutant sites.

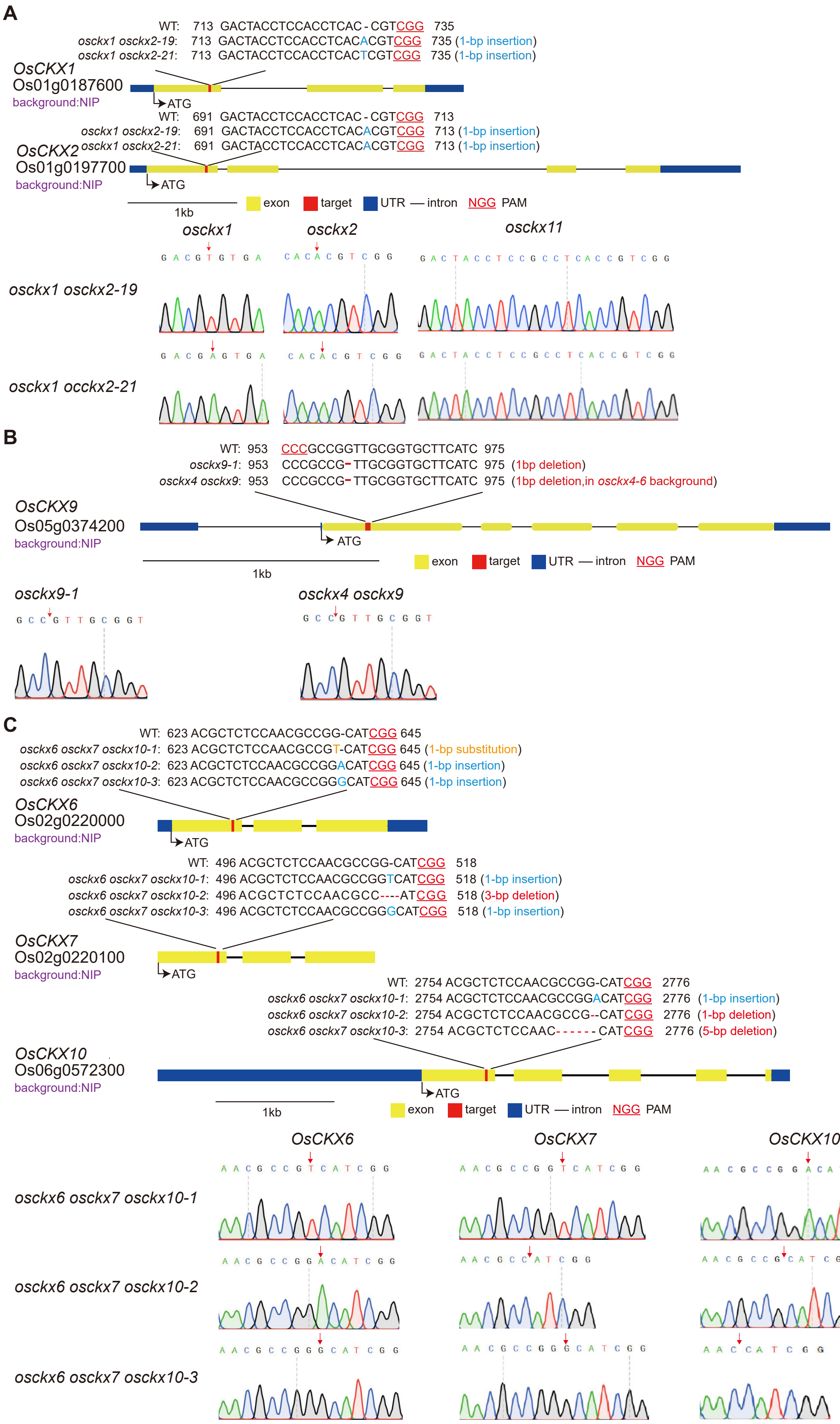


**Supplemental Figure S3.** Mutation details and chromatograms of double and triple mutants. A, Mutation details of the coding sequences and chromatograms in the *OsCKX1, OsCKX2* of *osckx1 osckx2* mutants used in research; B, Mutation details of the coding sequences and chromatograms in the *OsCKX9* of *osckx9* and *osckx4 osckx9* mutants used in research; C, Mutation details of the coding sequences and chromatograms in the *OsCKX6, OsCKX7* and *OsCKX10* of *osckx6 osckx7 osckx10* mutants used in research. The solid yellow, blue, and red boxes represent the exons, untranslated regions, and target sequences, respectively. The introns are shown as solid lines, while protospacer adjacent motif (PAM) sequences are shown in red color and underlined with red. Red arrows in chromatograms point to mutant sites.

A

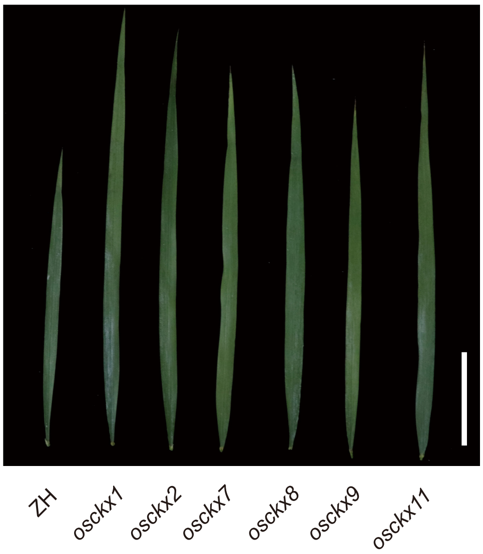


B

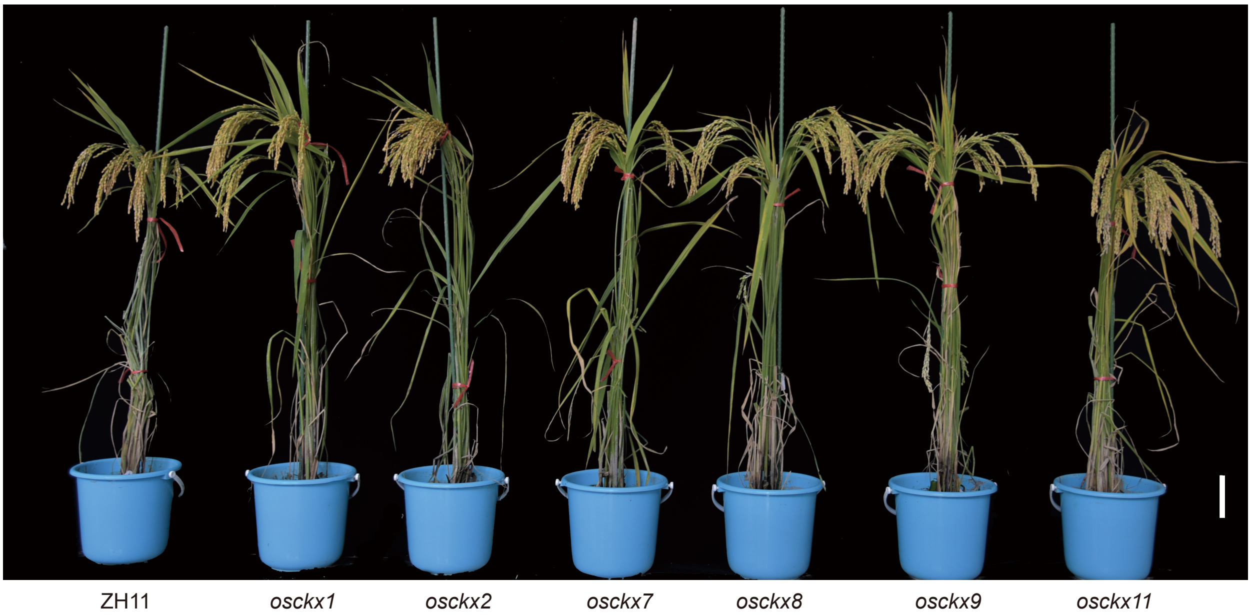


**Supplemental Figure S4.** The *osckx* mutants of Zhonghua 11 from the 2020 field experiment. A, Images of the flag leaves of *osckx* plants; B, images of the *osckx* plants at the ripening stage. Scale bar = 10 cm.

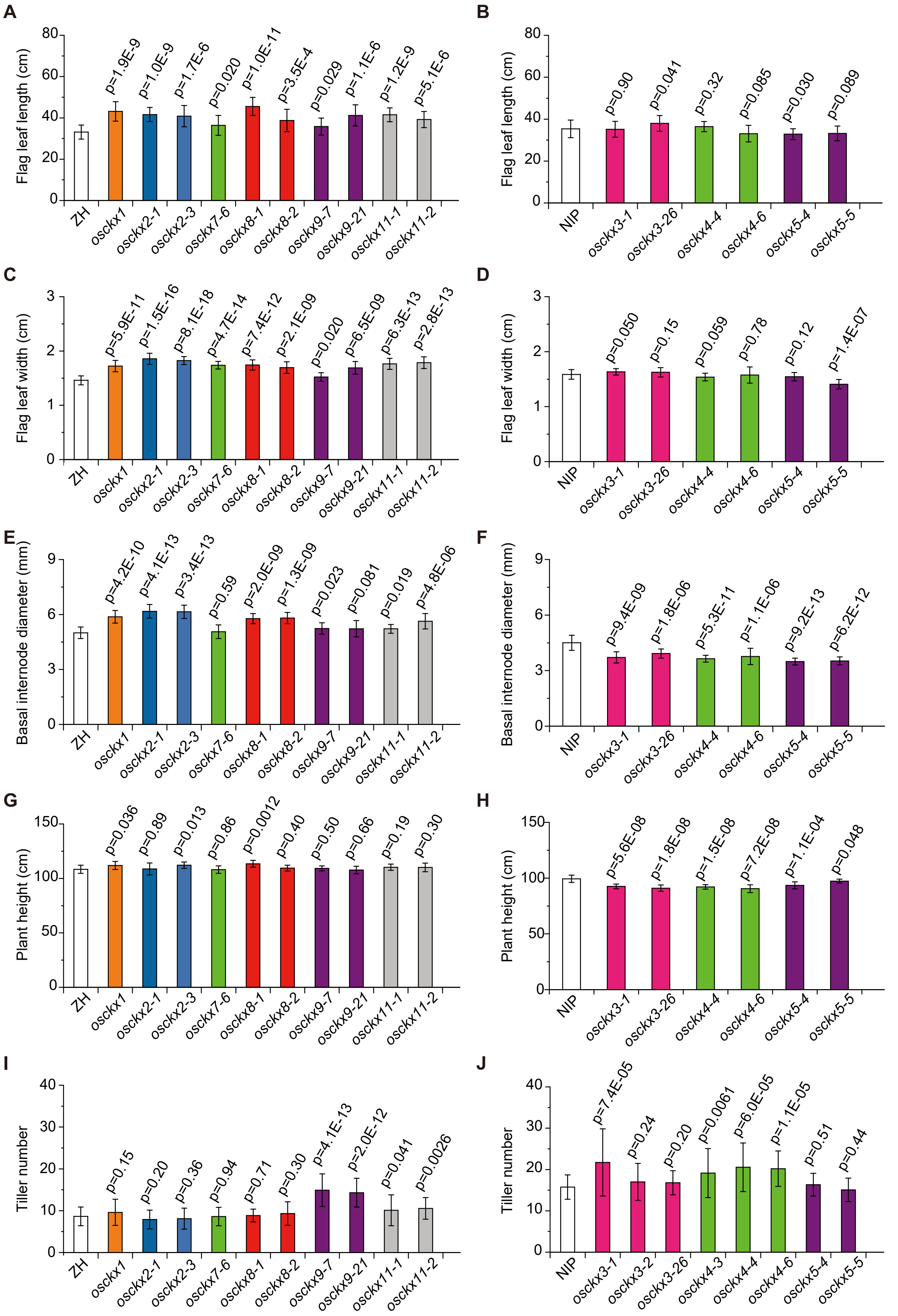


**Supplemental Figure S5.** Phenotypic characterization of the vegetative organs in *osckx* mutants from the 2020 field experiment. A,B, Flag leaf length of each mutant line for Zhonghua (ZH) (A) and Nipponbare (NIP) (B) background; C,D, Flag leaf width of each mutant line for ZH (C) and NIP (D) background; E,F, Basal internode diameter of each mutant line for ZH (E) and NIP (F) background; G,H, plant height of each mutant line for ZH (G) and NIP (H) background; I,J, tiller number of each mutation line at 69 days after sowing (reproductive stage) for ZH (I) and NIP (J) background. Data are shown as means ± SDs (n = 20); numbers above the error bars indicate the level of statistical significance from their background control by independent-samples *t*-test.

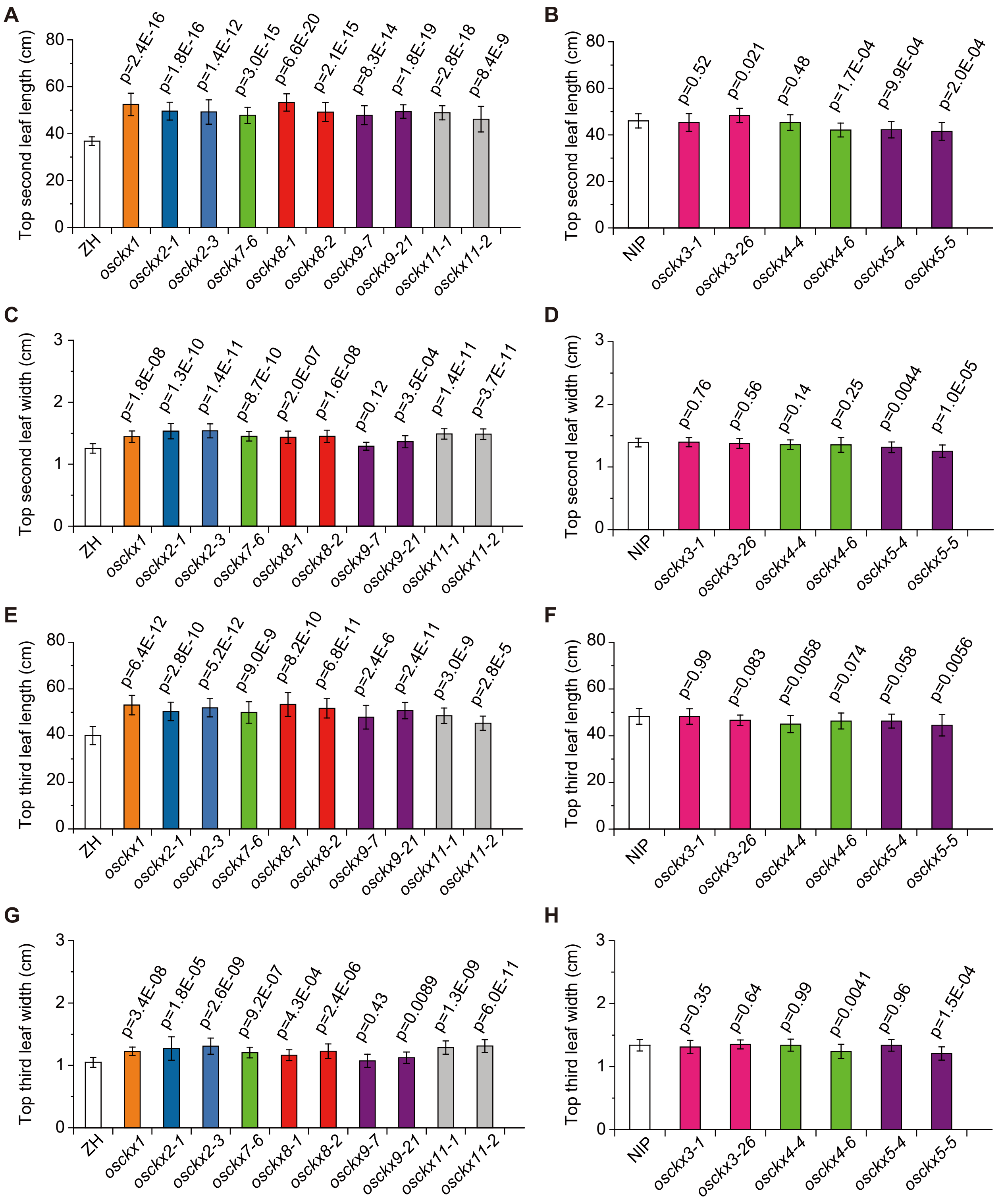


**Supplemental Figure S6.** Phenotypic characterization of the second and third top leaves in *osckx* mutants from the 2020 field experiment. A,B, Second top leaf length of each mutant line for Zhonghua (ZH) (A) and Nipponbare (NIP) (B) background; C,D, second top leaf width of each mutant line for ZH (C) and NIP (D) background; E,F, third top leaf length of each mutant line for ZH (E) and NIP (F) background; G,H, third top leaf width of each mutant line for ZH (G) and NIP (H) background. Data are shown as means ± SDs (n = 20); numbers above the error bars indicate the level of statistical significance from their background control by independent-samples *t*-test.

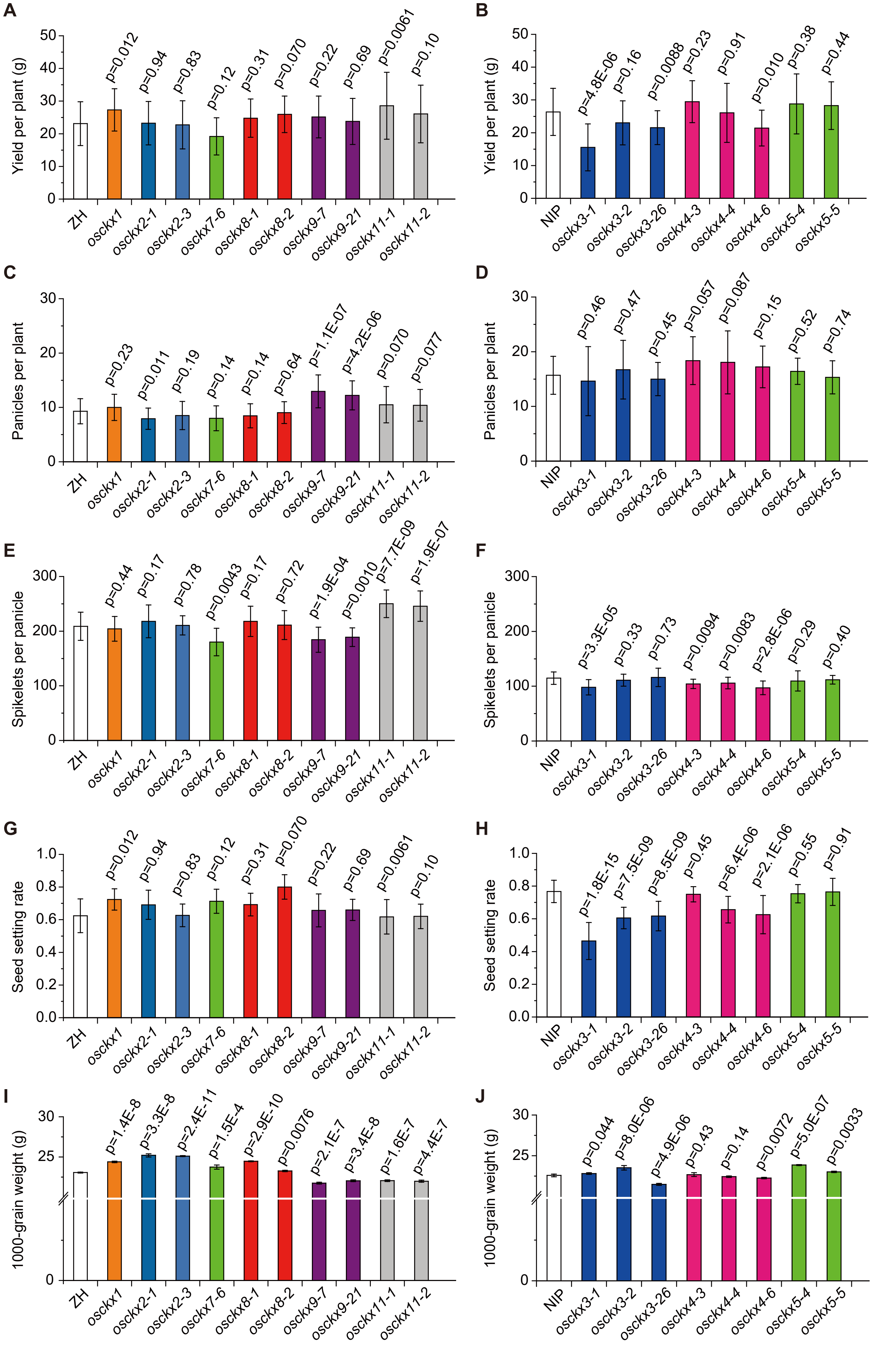


**Supplemental Figure S7.** Phenotypic characterization of yield-related phenotypes in *osckx* mutants from the 2020 field experiment. A,B, Yield per plant of each mutant line for Zhonghua (ZH) (A) and Nipponbare (NIP) (B) background; C,D, panicles per plant of each mutant line for ZH (C) and NIP (D) background; E,F, spikelets per panicle of each mutant line for ZH (E) and NIP (F) background; G,H, seed setting rate of each mutant line for ZH (G) and NIP (H) background; I,J, 1000-grain weight of each mutant line for ZH (I) and NIP (J) background. Data are shown as means ± SDs (n = 20 for A–H; n = 5 for I and J); numbers above the error bars indicate the level of statistical significance from their background control by independent-samples *t*-test.

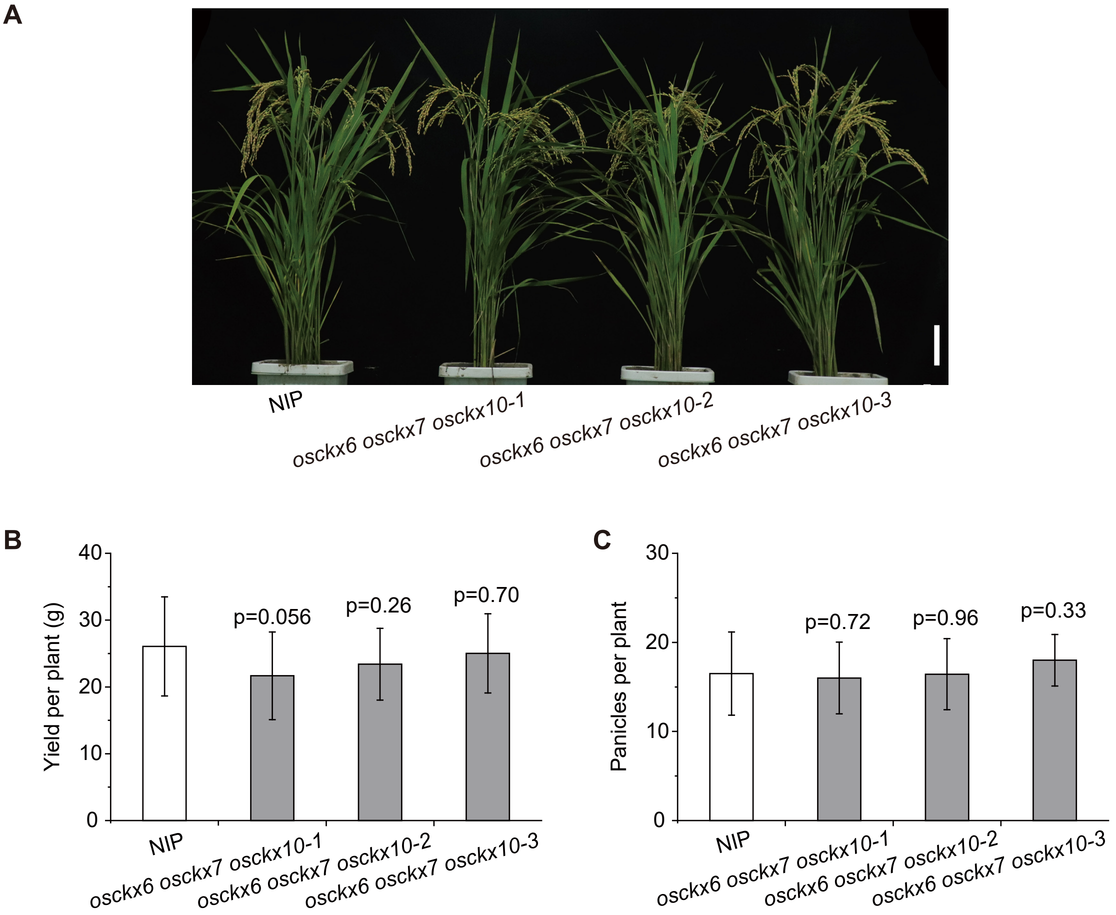


**Supplemental Figure S8.** The phenotypic characterization of the yield and panicles per plant in Nipponbare (NIP) and *osckx6 osckx7 osckx10* plants. A, Phenotypic features of Nipponbare (NIP) and *osckx6 osckx7 osckx10* plants at the mature stage. The image was digitally extracted and scaled for comparison (scale bar = 10 cm); B,C, Measurement of yield per plant (B) and panicles per plant (C) of NIP and *osckx6 osckx7 osckx10* (n=15) plants. The P-values indicate the level of statistical significance between NIP and *osckx6 osckx7 osckx10* determined by Student's *t*-test.

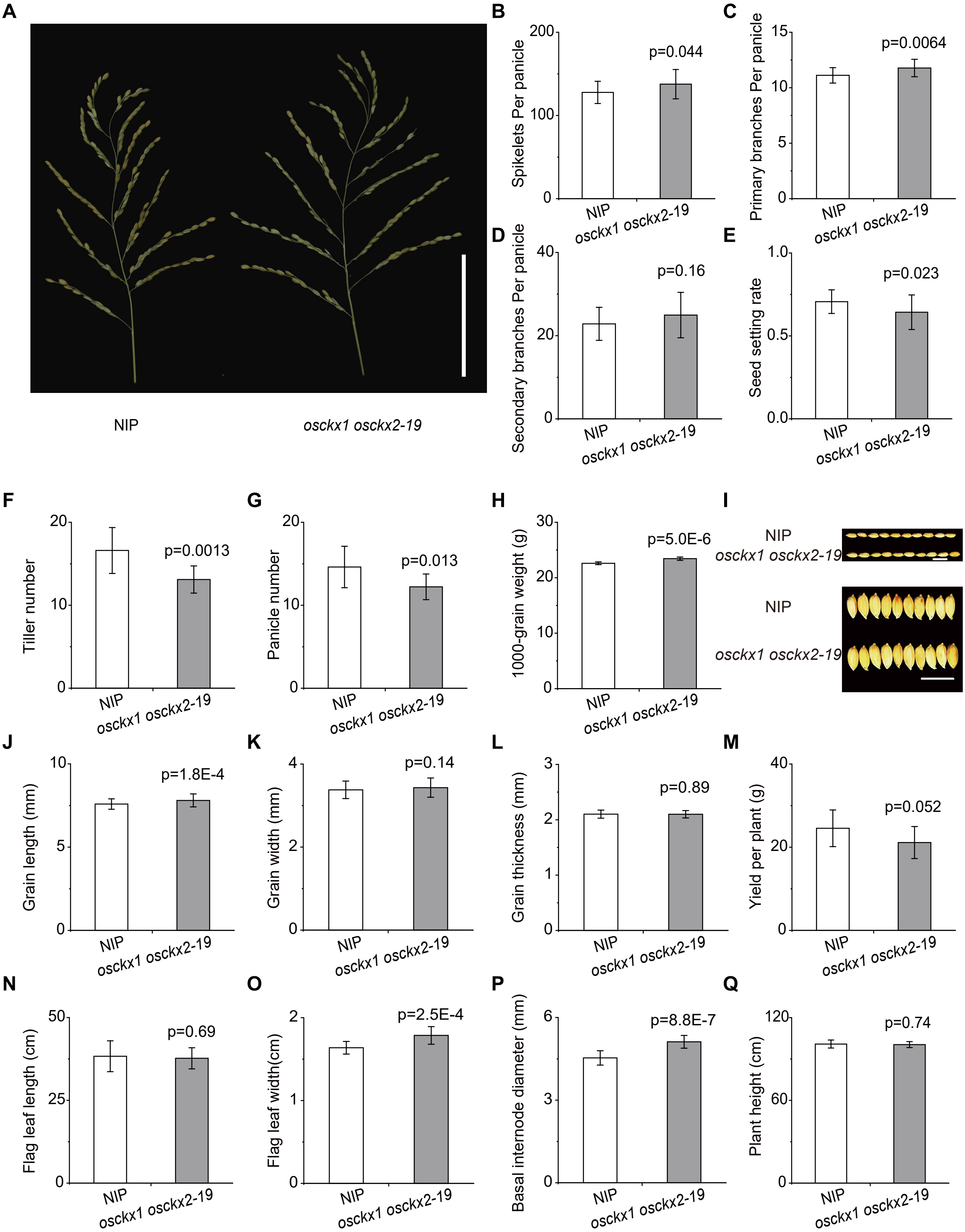


**Supplemental Figure S9.** Phenotypic characterization of Nipponbare (NIP) and *osckx1 osckx2-19* mutant plants in 2020. A, Panicle phenotypes of NIP and *osckx1 osckx2-19* plants. The image was digitally extracted and scaled for comparison (scale bar = 10 cm); B-H, Measurement of the spikelets per panicle (B), primary branches per panicle (C), secondary branches per panicle (D), seed setting rate (E), tiller number at the vegetative stage 70 days after sowing (F), panicle number at reproductive stage (G), 1,000-grain weight (H) between NIP (n > 20 for B–G, n = 5 for H) and *osckx1 osckx2-19* (n > 10 for B–G, n = 5 for H) plants; I, Grain phenotypes of NIP and *osckx1 osckx2-19* plants. The image was digitally extracted and scaled for comparison (scale bar = 1 cm); J-Q, Measurement of the grain length (J), grain width (K), and grain thickness (L), yield per plant (M), length of flag leaves (N), width of flag leaves (O), basal internode diameter (P), and plant height (Q) of NIP (n > 50 for J–L , n > 20 for M–Q) and *osckx1 osckx2-19* ((n > 50 for J–L, n > 10 for M–Q) plants. The P-values indicate the level of statistical significance between NIP and *osckx1 osckx2-19* determined by Student's *t*-test.

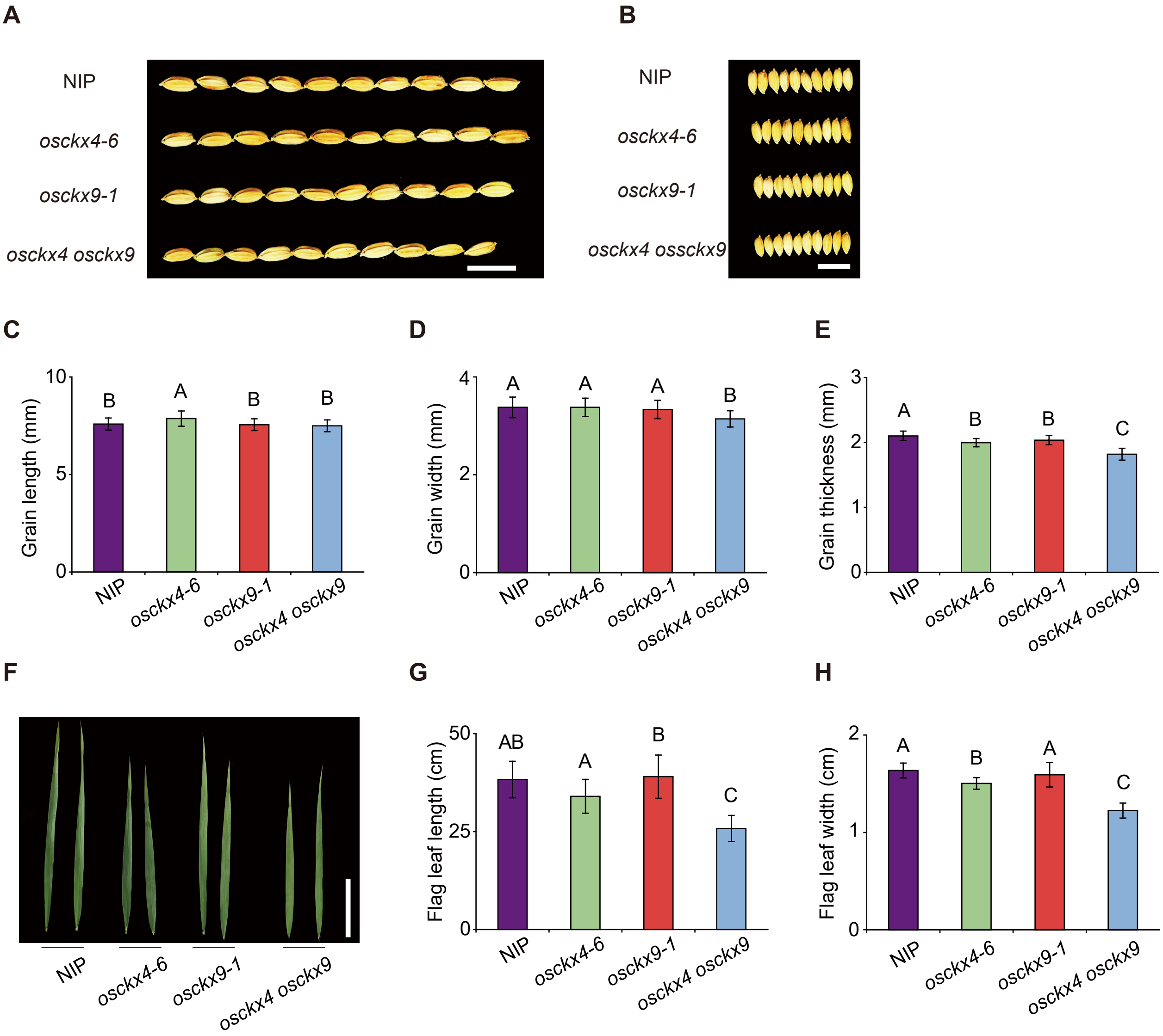


**Supplemental Figure S10.** Phenotypic characterization of the flag leaves and seeds of Nipponbare (NIP), *osckx4*, *osckx9*, and *osckx4 osckx9* plants. A,B, grain length (A) and width (B) phenotypic features of NIP, *osckx4-6*, *osckx9-1*, and *osckx4 osckx9*; images shown were digitally extracted and scaled for comparison; scale bar = 1 cm; C–E, measurement of grain length (C), grain width (D), and grain thickness (E) of NIP, *osckx4-6*, *osckx9-1*, and *osckx4 osckx9*; n > 50 for each index; F, phenotypic features of NIP, *osckx4-6*, *osckx9-1*, and *osckx4 osckx9* flag leaves; images shown were digitally extracted and scaled for comparison. Scale bar = 10 cm. G,H, measurement of flag leaf width (G) and leaf length (H); data are shown as means ± SDs (n = 15). Different capital letters indicate the level of statistical significance ($P < 0.01$) as determined by one-way ANOVA and shortest significant range mean-separation test.

**A**

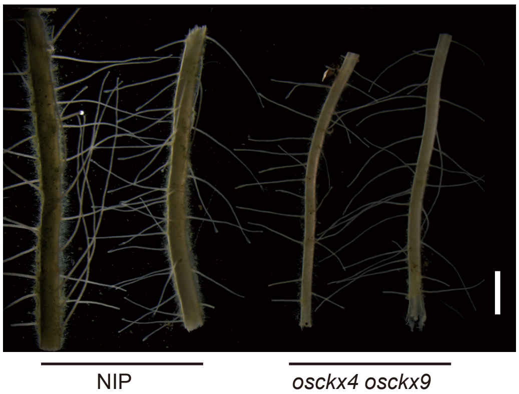


**B**

| root phenotypes | NIP | *osckx4 osckx9* |
|---|---|---|
| root length (cm) | 17.2±1.3 | 12.3±1.1 ** |
| root diameter (mm) | 0.77±0.12 | 0.51±0.06 ** |
| crown root number | 52.9±6.7 | 40.3±5.3 ** |

**Supplemental Figure S11.** Phenotypic characterization of the leaf and root systems in wild-type and *osckx4 osckx9* plants. A, Phenotypic features of Nipponbare (NIP) and *osckx4 osckx9* crown roots; images shown were digitally extracted and scaled for comparison; scale bar = 1 mm; B, measurement of root length, root diameter, and crown root number of NIP and *osckx4 osckx9*; data are shown as means ± SDs ; asterisks indicate the level of statistical significance between NIP and *osckx4 osckx9* based on independent-samples *t*-test (**P < 0.01; n = 12).

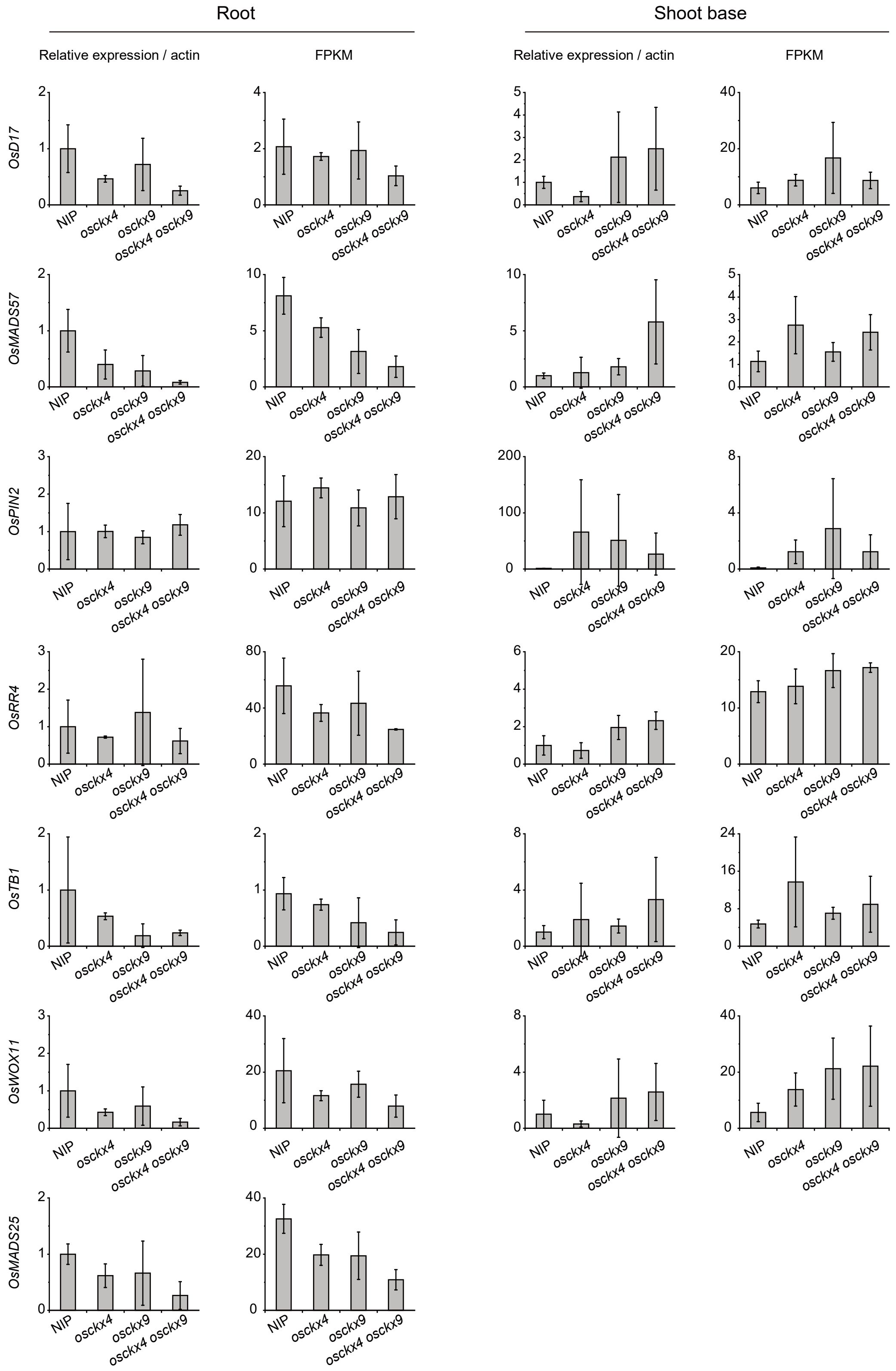


**Supplemental Figure S12.** Relative expression levels and FPKM values of *OsD17*, *OsRR4*, *OsWOX11*, *OsPIN2*, *OsTB1*, *OsMADS25*, and *OsMADS57* in the roots and shoot bases (BP) of Nipponbare (NIP), *osckx4*, *osckx9*, and *osckx4 osckx9*. Quantitative expressional changes in seven differentially expressed genes via RNA-seq and qRT-PCR. Results are presented relative to NIP. Both the FPKM value and qRT-PCR data represent means ± SD (n = 3). Total RNA was isolated from NIP or mutants' root or shoot base. β-Actin was used as a reference gene.

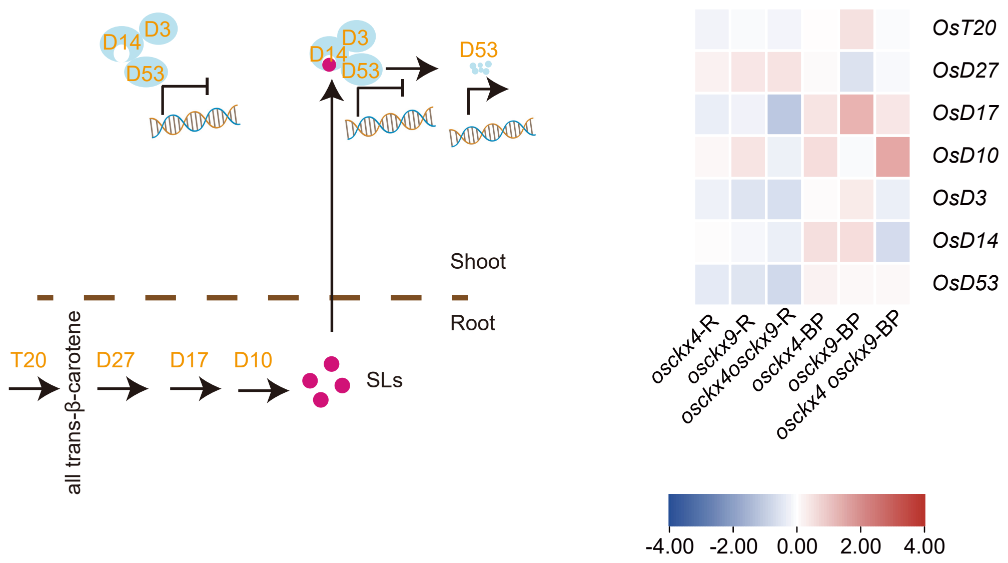


**Supplemental Figure S13.** OsCKX4 and OsCKX9 are not associated with strigolactone biosynthesis and signaling transduction. Differentially expressed genes (DEGs) related to strigolactone metabolism and signaling transduction in the root (R) and shoot base (BP) of different mutants compared with Nipponbare (NIP). The fold change was based on $\log_2$ ($FPKM_{mutant}/FPKM_{WT}$) values. * $Q < 0.05$, ** $Q < 0.01$.

**Supplemental Table S1** List of primers used in this study

| Primer | primer sequence | usage |
|---|---|---|
| OsCKX1-ZH mF | ACTACCTCCACCTCACCGT | |
| OsCKX1-ZH/NIP mR | CGAGGTTTGTATGTGTGAGG | |
| OsCKX1-NIP mF | GCAAGAGCACAACACCCTTA | |
| OsCKX2-ZH mF | CGGCGTCGGAGAAGAAGGAT | |
| OsCKX2-ZH mR | GCAGGTGTCGGGGAGATGGT | |
| OsCKX2-NIP mF | CTCGCCGTGTCAGTGGAG | |
| OsCKX2-NIP mR | CGGCGCGTGTCTTAGTAGAT | |
| OsCKX3-NIP mF | ATCCAGTGACTTTGGTAGAATC | |
| OsCKX3-NIP mR | TGTCAAGCAAAGAGGAACAGA | |
| OsCKX4-NIP mF | GATCGTCGTCAAGATGGAG | |
| OsCKX4-NIP mR | CTCCCTGCATTTGGTAGTTC | |
| OsCKX5-NIP mF | GGATAACTGTCGTGTGTGCTT | Genotyping |
| OsCKX5-NIP mR | TGGTCGTGGACATGAGTCAC | |
| OsCKX7-ZH mF | TGCTCAAGGGACAAGGACTC | |
| OsCKX7-ZH mR | TTCTTCCTCCCTCACACGAT | |
| OsCKX8-ZH mF | TCTATGAGAATGGAAGGGTC | |
| OsCKX8-ZH mR | GTCTTGCTGGAAATCGTG | |
| OsCKX9-ZH mF | CCAGAGTAACACAATGAGGG | |
| OsCKX9-ZH mR | ATAACATCACCTCTTCCTGC | |
| OsCKX9-NIP mF | AGCTGCTCCTGTTACTTGCC | |
| OsCKX9-NIP mR | CCAGGGTTCACCCTCATTGT | |
| OsCKX11-ZH/NIP mF | CGAGGACGGCGAAGAAGAGG | |
| OsCKX11-ZH/NIP mR | CGGCGAGCCAAGATGATGC | |
| OsCKX1-ZH-sgF | GGCAGGCGGGCAGACTTACCGGCA | |
| OsCKX1-ZH-sgR | AAACTGCCGGTAAGTCTGCCCGCC | |
| OsCKX2-ZH-sgF | GGCAGGTCCACCTGAACCAGGGCC | |
| OsCKX2-ZH-sgR | AAACGGCCCTGGTTCAGGTGGACC | |
| OsCKX7-ZH-sgF | GGCAGCCGGCTTCGACTACGTCGA | |
| OsCKX7-ZH-sgR | AAACTCGACGTAGTCGAAGCCGGC | |
| OsCKX8-ZH-sgF | GGCATCCCAAGGTACTAACTCTAC | |
| OsCKX8-ZH-sgR | AAACGTAGAGTTAGTACCTTGGGA | |
| OsCKX9-ZH-sgF | GGCAAGTGGGCAGACCTTCCGGCA | |
| OsCKX9-ZH-sgR | AAACTGCCGGAAGGTCTGCCCACT | Construct mutants |
| OsCKX11-ZH-sgF | GGCAGCGAGAAGTTCGCCGACGTC | |
| OsCKX11-ZH-sgR | AAACGACGTCGGCGAACTTCTCGC | |
| OsCKX3-NIP-sgF | GGCAGGACAGGCACAGGCCCTTGA | |
| OsCKX3-NIP-sgR | AAACTCAAGGGCCTGTGCCTGTCC | |
| OsCKX4-NIP-sgF1 | GGCAGGGTCATGGACCGACTACCT | |
| OsCKX4-NIP-sgR1 | AAACAGGTAGTCGGTCCATGACCT | |
| OsCKX4-NIP-sgF2 | GGCAGGTGGCACCTTGTCGAATGC | |
| OsCKX4-NIP-sgR2 | AAACGCATTCGACAAGGTGCCACC | |
| OsCKX5-NIP-sgF | GGCAGTATAAGTGGGCAAGCTTTC | |

| | | |
|---|---|---|
| OsCKX5-NIP-sgR | AAACGAAAGCTTGCCCACTTATAC | |
| OsCKX9-NIP-sgF | GGCAGATGAAGCACCGCAACCGGC | |
| OsCKX9-NIP-sgR | AAACGCCGGTTGCGGTGCTTCATC | |
| OsCKX1/2-NIP-sgF | GGCAGACTACCTCCACCTCACCGT | |
| OsCKX1/2-NIP-sgR | AAACACGGTGAGGTGGAGGTAGTC | |
| CKX1proPstF | CTGCAGTAACCCCTCCTAACCATCAACC | |
| CKX1proBamHR | GGATCCTAATAAGGGTGTTGTGCTCTTGC | |
| CKX2proSalF | GTCGACCACCGTAGCTATAGAGATTCAT | |
| CKX2proSalR | GTCGACTATCAATCAATCAATCGATCGGT | |
| CKX2terSac | GAGCTCAATGACACATGTATGCAAATGCAT | |
| CKX2terEcoR | GAATTCTCCCCCTGTCTCATAAAAAACG | |
| CKX3proSalF | GTCGACAATCAGCACAAAAGAAACAGGG | |
| CKX3proBamHR | GGATCCAAGTTGAGGGGATGAAGAAAAGG | |
| CKX4proPmeF | GTTTAAACGTCTGCGGAAAAATGTGTATGTG | Construct GUS vectors |
| CKX4proXbaR | TCTAGAGTGAGTGAGAGTTGTTGGGGG | |
| CKX5proPmeF | GTTTAAACCTCCTCCCATCTATCCACATCCACTATC | |
| CKX5proSalR | GGATCCAAGGAGGAGGAGGAGAGATGGCGATGT | |
| CKX8proPmeF | GTTTAAACAAGGGGCGTGTGAAAAGGTGCGAG | |
| CKX8proHindR | AAGCTTGTCGTGCTGCTCTCTGCTTGAT | |
| CKX9proSalF | GTCGACCAATACCCAACCACGATAAGAC | |
| CKX9proBamHR | GGATCCTGGCAATGTTTTGAGAGGAAGAC | |
| CKX11proXbaF | TCTAGAGTTGGAGGATTGTTGAGTTTGGC | |
| CKX11proBamHR | GGATCCTGCGTTTCTTGTTTCTCTGTCTC | |
| T20rtF | ATCACTAGACATCCACAGA | |
| T20rtR | CCTCCTATCACCATTCCA | |
| D27rtF | TGCTGCATTCACCACGATA | |
| D27rtR | ACCAACACAATTTGTACTTTCCA | |
| D17rtF | ATCCCACTTCACTTTCTACGAG | |
| D17rtR | CTGTTGCCGAGGAGGATGTAG | |
| D14rtF | CGCCTCTCCCCGGTTCTTG | |
| D14rtR | ATGTTGAAGAGGGTGCGGCTG | |
| D3rtF | GAACACAACCGAACAACCCTGC | |
| D3rtR | GCAGAAATGAGCGGAGAGAGAAG | |
| TB1-rtF | CGACAGCGGCAGCTACTAC | q-RT PCR |
| TB1-rtR | GCGAATTGGCGTAGACGA | |
| MADS57rtF | GCAAGAAAGCCACAAGCAAC | |
| MADS57rtR | CTCTTTGCTGTGATTGGCTC | |
| WOX11rtF | ACTACTCGTGTCAACCTGCG | |
| WOX11rtR | TACTCGTTGGCTGGAAGAAG | |
| MADS25rtF | TACACCACAACAACAGGCAAC | |
| MADS25rtR | TCAAGGTCAATACACACACGT | |
| PIN2-rtF | TGTCAGATGCAGGGCTAGGA | |
| PIN2-rtR | TATCCCAAGAAGCACATAGT | |
| RR1rtF | CAAGGAAGAAGGCAAGGAGCAG | |

| | |
|---|---|
| RR1rtR | TCCTCTGGGGCTCGTTTTCC |
| RR2rtF | CGGAGATGACAGGATACGAT |
| RR2rtR | CGCTGGACATCGTTCATCTT |
| RR3rtF | CGAGTTACGACGGTGGATAG |
| RR3rtR | CCGCCGACTCCTTGACCCTCT |
| RR4rtF | TCTTCTGAGAATGTGCCTGCAA |
| RR4rtR | TGTGTGGCGGCTTATCTGGTG |
| RR5rtF | GGAAGAGGGCATTGGAGCTG |
| RR5rtR | GCACCTGTTGATCCTTGTAG |
| RR6rtF | GTCCCCAACGTCAACATGATC |
| RR6rtR | TGAGACGATTCCTTGACGCG |
| RR7rtF | TGCTCAAGAAGATCAAGGAATCG |
| RR7rtR | GGCTGGTTATGCGGGAGATGT |
| RR9rtF | TGCTTTCTCCTTGTAGTCTCTTCTG |
| RR9rtR | CGGAATCAACAGTGGTAAC |
| RR10rtF | CTTTCTCCTTGTAGCCTCTTCCTT |
| RR10rtR | TCCCCGAATCAACAGTGGTT |
| RR11rtF | GAAGAAAGTCAAGGAGTCAT |
| RR11rtR | TCACCAAGAAATCCTCCGCG |
| actin-rtF | CAATCGTGAGAAGATGACCC |
| actin-rtR | GTCCATCAGGAAGCTCGTAGC |

**Supplemental Table S2** Agricultural traits in field in 2019

| mutant lines | tiller number | flag leaf length (cm) | flag leaf width (cm) | panicles each cave | spikelets per panicle | panicle weight (g) | yield per cave (g) |
|---|---|---|---|---|---|---|---|
| ZH | 22.2 ± 3.8 | 28.8 ± 3.2 | 1.28 ± 0.09 | 21.4 ± 4.6 | 174.3 ± 19.9 | 2.85 ± 0.32 | 41.73 ± 8.74 |
| *osckx1* | 19.4 ± 3.3 * | 35.6 ± 3.8 ** | 1.47 ± 0.07 ** | 18.9 ± 2.9 | 177.3 ± 23.8 | 3.09 ± 0.47 * | 37.20 ± 9.18 |
| *osckx2-1* | 16.3 ± 3.9 ** | 32.3 ± 4.2 ** | 1.62 ± 0.12 ** | 16.0 ± 3.3 ** | 216.7 ± 22.3 ** | 4.22 ± 0.43 ** | 43.40 ± 10.94 |
| *osckx2-3* | 17.2 ± 3.5 ** | 36.7 ± 5.4 ** | 1.53 ± 0.09 ** | 17.2 ± 3.6 * | 183.4 ± 20.6 | 3.41 ± 0.37 ** | 40.85 ± 8.17 |
| *osckx9-7* | 27.8 ± 4.6 ** | 29.8 ± 2.7 | 1.35 ± 0.08 * | 27.2 ± 4.8 ** | 150.1 ± 10.4 ** | 2.55 ± 0.25 ** | 43.29 ± 9.87 |
| *osckx9-21* | 27.2 ± 3.8 ** | 31.5 ± 4.2 | 1.38 ± 0.07 ** | 23.5 ± 2.9 | 153.7 ± 18.6 ** | 2.52 ± 0.41 ** | 37.46 ± 9.74 |

* for significantly different from ZH at $P < 0.05$, ** for significantly different from ZH at $P < 0.01$ (Student's t-test).

**Table S3** Tiller numbers in pot in 2019

| mutant lines | tiller number | |
|---|---|---|
| ZH | 9.8±1.9 | |
| *osckx1* | 9.8±1.5 | |
| *osckx2-1* | 6.5±1.4 | ** |
| *osckx2-3* | 8.8±1.9 | |
| *osckx7-3* | 8.2±1.4 | ** |
| *osckx7-6* | 7.8±1.8 | ** |
| *osckx8-1* | 7.4±1.7 | ** |
| *osckx8-2* | 8.9±1.6 | |
| *osckx9-7* | 11.8±2.5 | ** |
| *osckx9-21* | 13.1±3.1 | ** |
| *osckx11-1* | 10.2±1.4 | |
| *osckx11-2* | 9.9±1.6 | |
| NIP | 13.4±2.6 | |
| *osckx3-26* | 13.7±3.1 | |
| *osckx4-5* | 15.4±2.4 | * |
| *osckx4-6* | 17.4±3.2 | ** |

* for significantly different from ZH at P < 0.05, ** for significantly different from ZH at P < 0.01 (Student's t-test).
